# GTPase-dependent cyclic flexibility transitions drive the two-component EEA1-Rab5 molecular motor

**DOI:** 10.1101/2022.07.18.500463

**Authors:** Anupam Singh, Joan Antoni Soler, Janelle Lauer, Stephan W. Grill, Marcus Jahnel, Marino Zerial, Shashi Thutupalli

## Abstract

The recognition of vesicles by their correct target compartment depends on the pairing of small GTPases with effector proteins (1–3). Whereas ATPases can cyclically convert the free energy of ATP hydrolysis into mechanical work, the function of small GTPases is predominantly associated with signal transduction processes, and their role in mechano-transduction is less established (4–6). However, binding of the GTPase Rab5 to the long coiled-coil tethering protein EEA1 on an early endosome induces a rigidity transition resulting in a large conformational change in EEA1 from a rigid and extended to a flexible and collapsed state. This entropic collapse of EEA1 gives rise to an effective force that can pull tethered membranes closer (7). It currently remains unclear if EEA1 can return from the collapsed to the extended conformation without the aid of chaperones. Here, we use fluorescence correlation spectroscopy to reveal that EEA1 in bulk solution can undergo multiple flexibility transition cycles that are associated with the binding and release of Rab5(GTP) and Rab5(GDP). Using semi-flexible polymer theory we provide evidence that the cyclic transitioning of Rab5-EEA1 between extended and collapsed conformations is driven by the energetics of Rab5 binding/unbinding and GTP hydrolysis. Cyclic flexibility transitions represent a complete mechanical work cycle that is able to perform up to 20 k_B_T of mechanical work per cycle, against an opposing force. Hence, Rab5 and EEA1 constitute a two-component molecular motor driven by the chemical energy derived from GTP hydrolysis by Rab5. We conclude that coiled-coil tethering proteins and their small GTPase partners can have active mechanical roles in membrane trafficking.

## Introduction

Intracellular traffic involves a complex choreography of mechanical and chemical steps, from the formation of a vesicle, and its movement along cytoskeletal tracks, up to the tethering and fusion with its appropriate target membrane compartment (1, 2, 8). Molecular motors that cyclically transduce chemical energy (ATP) to transport vesicles over long distances are a canonical example of the coupling of chemistry with mechanics. However, such a coupling is less understood for membrane tethering, driven by the pairing of small GTPases with either multi-subunit complexes (9, 10) or long dimeric coiled-coil tether molecules (11). Recently, conformational changes of the early endosomal tether EEA1, caused by its binding to the small GTPase Rab5, have been shown to result in mechanical forces pulling membranes in close proximity to each other (7). The prevalence of dimeric coiled-coil motifs in tethering molecules suggests that these long molecules can play generic mechanical roles in regulating and overcoming distance barriers that physically separate membranes (7, 12), thus facilitating fusion. EEA1 is a coiled-coil dimeric molecule with a contour length of 222 ± 26 nm (7) and binds to the small GTPase in its “active” GTP-bound form, Rab5(GTP), but not in its “inactive” GDP-bound form, Rab5(GDP) (Fig. 1**a**). Upon binding to Rab5(GTP) via a Rab5 binding domain located near the N-terminus, EEA1 undergoes a general change in conformation, from a more rigid, “extended” state to a more flexible “collapsed” state (7) (Fig. 1**a**). The flexibility transition of EEA1 causes it to collapse, *i*.*e*. adopt lower end-to-end distance configurations, due to entropic reasons thus generating an effective force than can bring tethered membranes closer (Fig. 1**a**). It is noteworthy that such entropic collapse associated with a flexibility transition along the length of the molecule could be shared by other coiled-coil tethers such as GCC185, whose flexibility is mediated by a local unwinding of specific sequences in the central region (8).

**Fig. 1.**
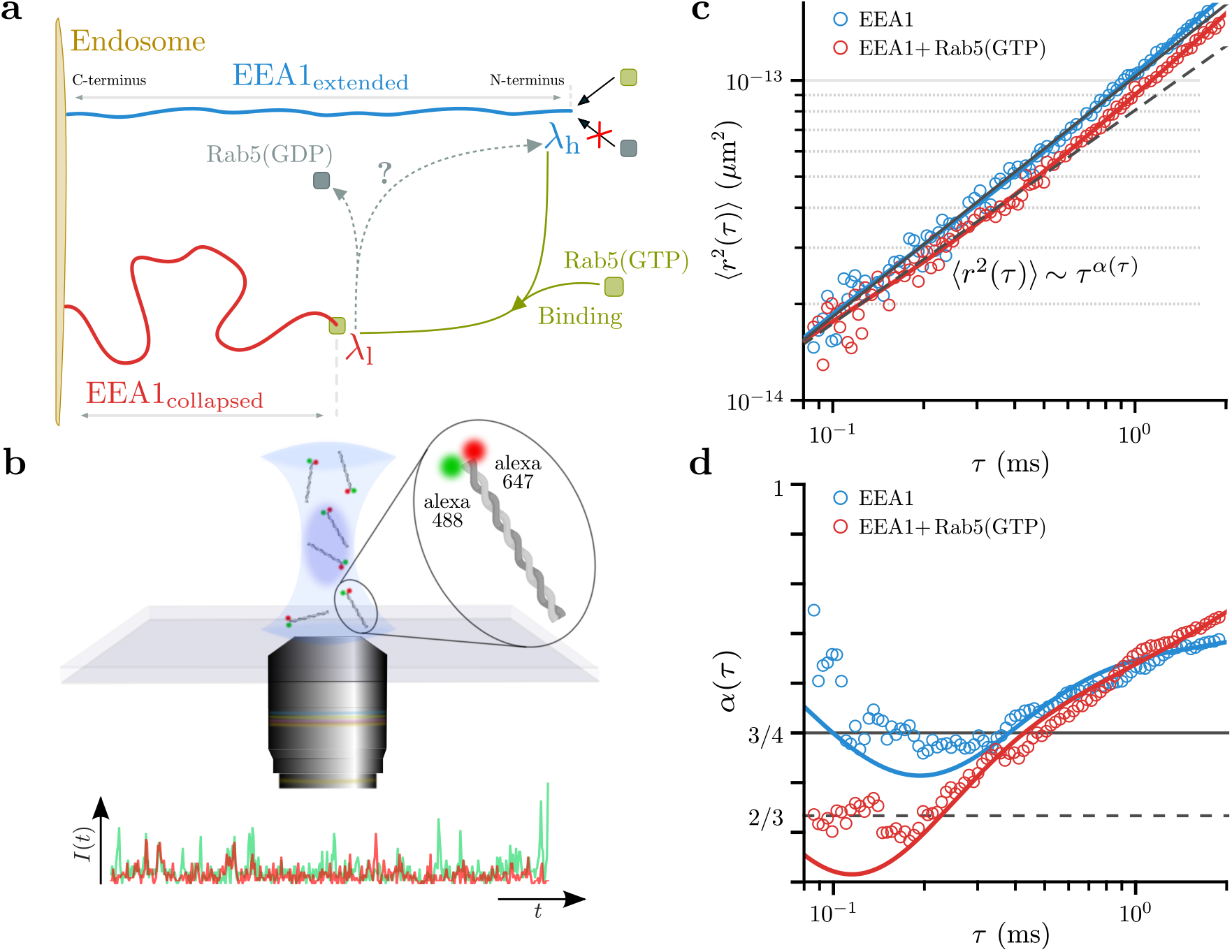
EEA1 undergoes a Rab5-dependent flexibility transition that can be observed in FCS experiments. Binding of the Rab5(GTP) to EEA1 triggers a transition of the EEA1 molecule from a more rigid “extended” state to a more flexible “collapsed” state. **(a)** Sketch depicting the collapsed/extended states of EEA1 upon binding/unbinding to the active/inactive forms of Rab5. **(b)** A doubly tagged (Alexa 488 and 647) EEA1 molecule is used in dcFCCS experimental setup to measure the dynamics of one end of the EEA1 molecule in solution. The time-series of the fluorescence intensity fluctuations within the confocal volume are used to quantify the dynamics of the molecular motion. The dynamics in solution for EEA1 alone (blue) and when mixed with Rab5(GTP) (red) are quantified by the **(c)** mean square displacement (MSD) plotted over a lag time *τ* and **(d)** the local scaling exponent *α* of the MSD. The solid and dashed black lines represent scaling exponents of *α* = 3/4 and *α* = 2/3 respectively, corresponding to regimes where the persistence length of EEA1 is comparable to its contour length *i*.*e*. “extended” and smaller than the contour length i.e. “collapsed”.

A high flux of vesicles in the endosomal system (∼1000 clathrin coated vesicles per minute (13)) necessitates the recycling of the EEA1 molecules, following the collapse mechanism, for new rounds of vesicle tethering and fusion. Thus, after the collapse to a more flexible configuration, EEA1 must regain its extended conformation *i*.*e*. switch its flexibility back to a stiffer state. However, it remains unclear whether completing a cycle of EEA1 collapse and extension requires chaperones, or if the transition back to an extended configuration can be supported by the energy of the GTPase cycle alone (Fig. 1**a**). The latter scenario would make the system similar to molecular motors like kinesin or myosin (14), with the key difference that the hydrolysis cycle would not drive the cyclic movement of a lever arm to perform mechanical work against an opposing force (14), but instead supports a molecular flexibility transition that results in a possibly reversible entropic collapse. Here, we resolved this question by combining dual-color fluorescence cross-correlation spectroscopy (dcFCCS) (15–17) and semi-flexible polymer theory (18, 19) to measure and interpret the conformational dynamics of EEA1 upon interaction with its GTPase partner Rab5. We found that no external chaperones are required for the conformational cycling of EEA1. Further, this reversible process results in a mechanochemical work cycle establishing EEA1 and the GTPase Rab5 as a two component molecular motor system.

## Results

### EEA1 flexibility switches coupled to Rab5(GTP) are captured by the cross-over dynamics of semi-flexible polymers

The dynamics of a long molecule, such as EEA1, in solution – comprised of intra-molecular motions *i*.*e*. bending, rotation and also the center-of-mass displacement of the entire macromolecule – are similar to that of a randomly moving polymer. In addition to the effects of hydrodynamics due to the surrounding solvent, these dynamics are predominantly affected by the flexibility of the polymer, which is quantified by the persistence length, *λ*, in relation to its total length *i*.*e*. contour length *L* (20). Particularly, *λ* and *L* affect the cross-overs from the dynamics due to bending modes on shorter timescales to rotation and center-of-mass diffusion on longer timescales. Consequently, evaluation of the complex cross-over dynamics characterizes the stiff (*λ* ≳ *L*) or flexible state (*λ* << *L*) of the polymer. Following previous experiments and analysis based on semi-flexible polymer theory, these cross-overs can be inferred by measuring the mean-squared-displacement (MSD) of one end of the long polymer molecule using fluorescence correlation spectroscopy (FCS) (16, 21, 22).

In order to perform FCS measurements on EEA1, we fluorescently labelled one of its termini. Owing to its size, the timescales corresponding to the center-of-mass diffusion of EEA1 through an FCS confocal volume are on the order of a few milliseconds while the dynamics due to the internal molecular bending and rotational motions occur on relatively faster time scales *i*.*e*. 10s–100s of microseconds. Often, in a technique such as FCS, these fast dynamics can be masked due to the photo-physical processes of the fluorescent molecule that occur on similar timescales (17). Therefore, to mitigate these effects and to access the intra-molecular dynamics, we performed dual-colour fluorescence cross-correlation spectroscopy (dcFCCS) on EEA1 molecules that are simultaneously tagged at the same end with fluorescent molecules of two spectrally non-overlapping colours (Alexa 488 and 647) *i*.*e*. dual-labelled-EEA1 (Fig. 1**b** and Supplementary Information Sec. I and Sec. II). To achieve dual-labelling of EEA1, we took advantage of its dimeric nature and engineered its C-terminal sequence to add the recognition sequence for the *S. aureus* Sortase A (SrtA) enzyme (Extended Data Figure 1, Methods and Supplementary Information Sec. I.B) — the C-terminus is chosen specifically to avoid interference with the binding of Rab5(GTP) at the N-terminal end of EEA1 (23) (Fig. 1**a**). We then performed the SrtA reaction to obtain a population of EEA1 molecules consisting of monomers labelled at their C-terminal end with either of two fluorescent tags (Fig. 1**b** inset).

The FCS measurements were performed by the continuous recording of a time series of the fluorescence intensity fluctuations of the dual labelled EEA1 at a concentration of 100 nM (50 *µ*l total volume *i*.*e*. in a dilute regime) within a confocal volume of 0.285 *µ*m^3^ (Fig. 1**b**). The measurements were performed before and immediately after the addition of 2 *µ*M Rab5(GTP) to the same EEA1 sample, a concentration that ensures the binding of most of the EEA1 molecules to Rab5(GTP) (23). The time series data are used to compute the time cross-correlations across the two colours, *C*(*τ*) from which the mean-squared displacement ⟨*r*^2^(*τ*)⟩ of one end of the EEA1 with and without binding to Rab5(GTP) was extracted using a suitable mathematical transformation (Methods, more details are in the Supplementary Information Sec. II).

The MSD corresponding to EEA1 before (blue data points, Fig. 1**c**) and after binding to Rab5(GTP) (red data points, Fig. 1**c**) show a marked difference in scaling behaviours characterized by the scaling exponent *α*(*τ*), where ⟨*r*^2^(*τ*)⟩ ∼ *τ*^*α*^. At long enough time scales, due to the diffusive motion of the molecular center-of-mass, the MSD scales linearly with *τ i*.*e*. the exponent *α*(*τ*) ≈ 1 for both these EEA1 states. Strikingly, at intermediate timescales, corresponding to the dynamics due to internal polymer motions, this exponent is different for the unbound and Rab5(GTP) bound forms of EEA1 — whereas *α* ∼ 3/4 for the unbound EEA1, we find that it is significantly lowered, *i*.*e. α* ∼ 2/3, upon the addition and thereby binding of Rab5(GTP) to EEA1 (Fig. 1**d**). The scaling behaviour of *α* ∼ 2/3 is what is expected for long, flexible polymers *i*.*e. λ* << *L*, the so-called Zimm-scaling regime for semi-flexible polymers, while the exponent *α* ∼ 3/4 corresponds to the so-called rigid-rod limit *i*.*e. λ* ≳ *L* (16, 18, 21).

Given that the contour length of EEA1 remains mostly unchanged upon Rab5(GTP) binding (7), the measured change in the scaling exponent strongly suggests a reduction in the persistence length *λ* of EEA1 upon binding to Rab5(GTP). Using the theoretical analysis of Hinczewski *et al*., we extracted the persistence length, *λ*, of EEA1 in the bound and unbound states by fitting the MSD to

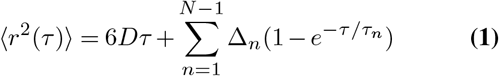

where the quantities *D*, Δ_*n*_ and *τ*_*n*_, *i*.*e*. the long-time center-of-mass diffusivity, length and time-scales corresponding to the rotation and bending motions, all depend on the persistence length, *λ* and the contour length, *L*. Indeed, from the analysis, we find that the persistence length of EEA1 undergoes a reduction upon Rab5(GTP) binding 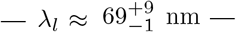 in comparison with the persistence length of free EEA1, 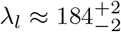 nm such a reduction is consistent with the stiffness transition previously characterized (7), the scaling behaviour that we have measured here highlights two key aspects: (i) the softening of EEA1 upon its binding to Rab5(GTP) likely occurs throughout its length *i*.*e*. EEA1 undergoes a global mechanical switch from a rigid-rod like polymer to a flexible one and (ii) a continuous measurement of the scaling opens a window into the long-term tracking of the EEA1 polymer mechanics.

### EEA1 flexibility transition is reversible without requiring additional components

We next asked whether the flexibility switch of EEA1 induced by Rab5(GTP) binding can be reversed and if it can occur cyclically without the aid of additional factors such as chaperones. In order to evaluate this, we tracked the long-term behaviour of a population of EEA1 molecules after the addition of 2 *µ*M Rab5(GTP), under the aforementioned experimental conditions. By measuring the changes in scaling exponent *α*, we found that the population of EEA1 molecules recovers to a rigid state similar to free EEA1 (Fig. 2**a**). The kinetics of this recovery, of the population of EEA1 molecules to their original state, are consistent with the intrinsic (low) bulk GTPase activity of Rab5 at 25°*C* (*k*_hydrolysis_ ≈ 5.5 × 10^−4^*s*^−1^) (24), suggesting that the recovery is coupled to the hydrolysis of GTP. The kinetics of Rab5 GTP hydrolysis (intrinsic rates) would imply that after 70 minutes, little (< 10%) active Rab5 remains in the solution (24).

**Fig. 2.**
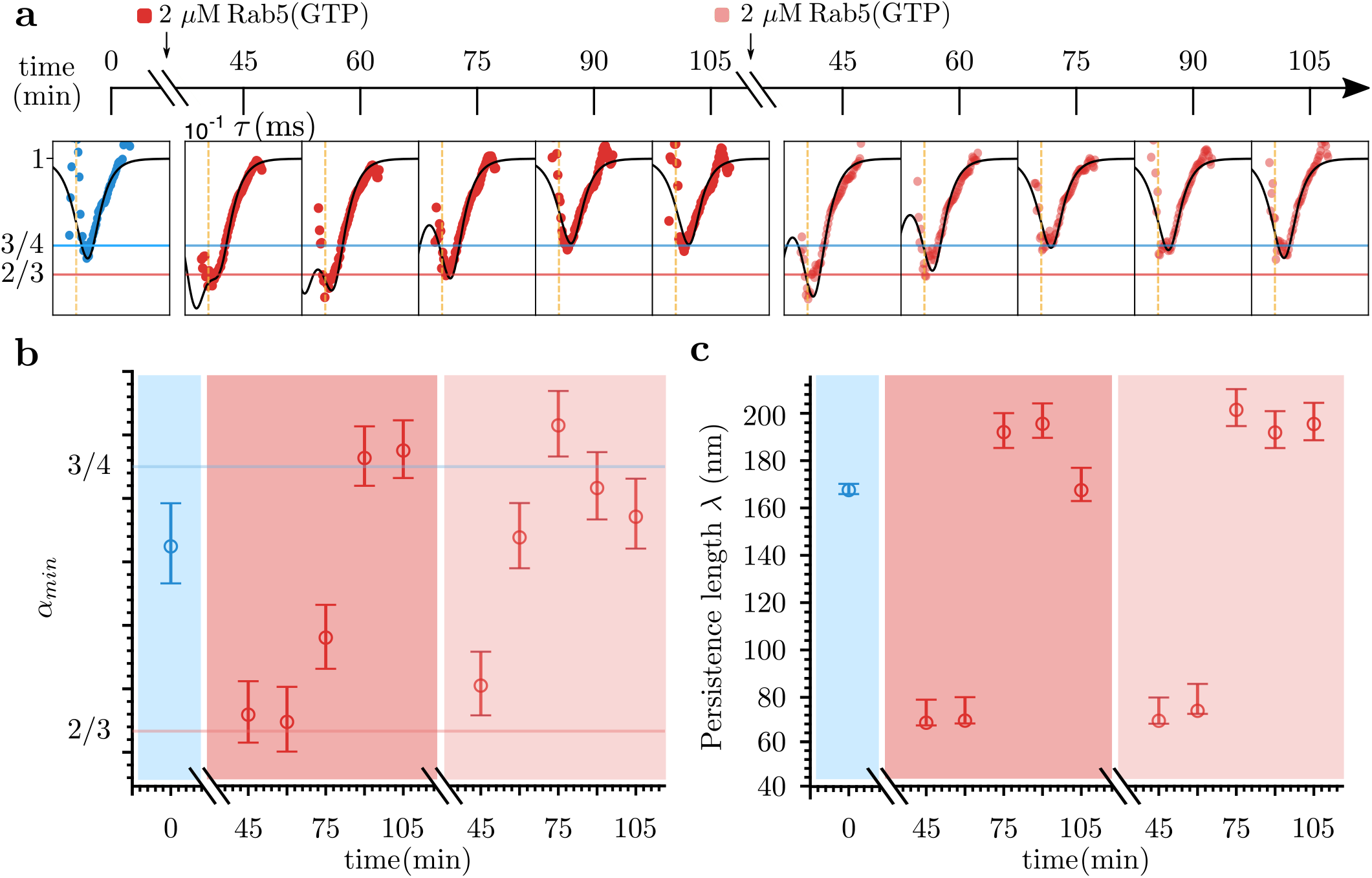
EEA1 and Rab5 spontaneously drive collapse-extension cycles without the assistance of additional components. **(a)** EEA1 alone (blue) in the extended state undergoes an entropic collapse (red) upon Rab5(GTP) addition followed by a recovery to the original extended state over time. A second round of Rab5(GTP) addition results in another EEA1 collapse-extension cycle. The filled colored circles corresponds to experimental data and solid black lines are fit to Eq. 1. The collapse-extension cycles are quantified from the changes over time in the **(b)** local slope minima *α*_*min*_ and **(c)** persistence length *λ*. For both cases, the corresponding peak value and the standard error of the peak (error bars) are calculated by bootstrapping (sample size = 10^5^). The quantified results in **(b)** and **(c)** show that the extended state of EEA1 (blue, *α*_*min*_ ≈ 3/4 and *λ ≈* 170 nm) undergoes a collapse (red, *α*_*min*_ ≈ 2/3 and *λ* ≈ 70 nm) upon binding to Rab5GTP, an over time population recovery to the original extended state (red, *α*_*min*_ ≈ 3/4 and *λ* ≈ 190*nm*), and a second collapse-extension cycle upon Rab5(GTP) addition. The blue and red solid lines correspond to the *α*_*min*_ = 3/4 (extended state) and *α*_*min*_ = 2/3 (collapsed state).

The ‘recovered’ state of EEA1 is similar in terms of its mechanical properties to the initial unbound EEA1, with a persistence length *λ* ≈ 190 ± 5 nm (Fig. 2**b,c** and Supplementary Information Sec. II.F). These results suggest that EEA1 has regained its extended conformation. If EEA1 has indeed been fully recycled to its initial extended state, it should be able to undergo new cycles of collapse and extension upon re-addition of Rab5(GTP). Such a recycling would not be possible if the EEA1 molecule were to enter an energetically proximate, yet inactive intermediate state that requires other input, such as the activity of chaperones, in order to be restored to its fully ‘active’ state. To test this, we measured a second cycle of entropic collapse by adding a fresh aliquot of 2 *µ*M active Rab5(GTP) to a solution of EEA1 that had undergone a cycle of collapse and re-extension. Remarkably, the recovered EEA1 collapsed once again into the flexible state and then recovered to the extended form (Fig. 2**a** second cycle). Note that the persistence length *λ* values are similar in both cycles. These results indicate that EEA1 can reversibly undergo multiple stiffness–flexibility transitions (Fig. 2**c**). These global flexibility transitions of EEA1 are triggered solely by the interaction with active Rab5(GTP) and do not require any additional factors, as a minimal system.

Finally, in all our experiments the MSD scaling exponent for the population of EEA1 upon recovery was measurably higher than that for free EEA1 *i*.*e*. we measured *α >* 3/4, concomitantly with a slight increase in effective persistence length (Fig. 2**b,c**). We speculated that this might be due to the presence of Rab5(GDP) in the solvent that does not bind to EEA1. To test this hypothesis, we performed separate experiments in which we added 2 *µ*M Rab5(GDP) to free EEA1. Consistent with our predictions, we found that the addition of Rab5(GDP) does not cause the EEA1 to transition to a flexible state. Strikingly however, we found that the scaling exponent for the EEA1 dynamics in the presence of the Rab5(GDP) is slightly higher than for free EEA1 *i*.*e. α >* 3/4 (Extended Data Figure 5 and Supplementary Information Sec. II.F). This increase in the scaling exponent is indeed consistent with what we observed in our recovery experiments, suggesting that Rab5(GDP) has a similar effect of increasing the scaling exponent. These data, together with the recovery kinetics, suggest that whereas Rab5(GTP) binding could account for the softening transition of EEA1, GTP hydrolysis and Rab5(GDP) unbinding may participate in the recovery of EEA1 back to the extended state.

Altogether, these measurements demonstrate that the long coiled-coil tether EEA1 can repeatedly undergo reversible flexibility transitions upon interaction with active Rab5 and that these transitions occur without the aid of external agents. These observations, combined with previous optical tweezer measurements of the force generated during the softening and subsequent collapse of EEA1 (7), suggest that EEA1 and Rab5 are involved in a mechanochemical work cycle *i*.*e*. they form a ‘two-component molecular motor’ system. By a two-component molecular motor we mean that its mechanical aspects (*i*.*e*. stiffness changes, force generation) and the chemical aspects (*i*.*e*. binding, GTP hydrolysis, phosphate release and unbinding) are attributed to two separate molecules — EEA1 and Rab5 respectively — that act in concert.

### A switchable semi-flexible polymer model for the EEA1–Rab5 molecular motor

To discuss the thermodynamics of the work cycle of EEA1 and Rab5, we consider a simplified mechanochemical model (Fig. 3). We assume that EEA1 is a semi-flexible polymer with a bending rigidity *κ* that is constant along the length of the molecule and that is related to a persistence length, *λ*, according to *λ* = *κ/*k_B_T. The bending rigidity is a mechanical quantity that includes various intra- and inter-molecular interactions, such as the pairing of heptad repeats in the EEA1 coiled coil, in a coarse-grained manner. The interaction of EEA1 with Rab5 leads to an instantaneous change in the bending rigidity *κ* that can be followed by a mechanical equilibration of the semi-flexible polymer via a change of its end-to-end distance.

**Fig. 3.**
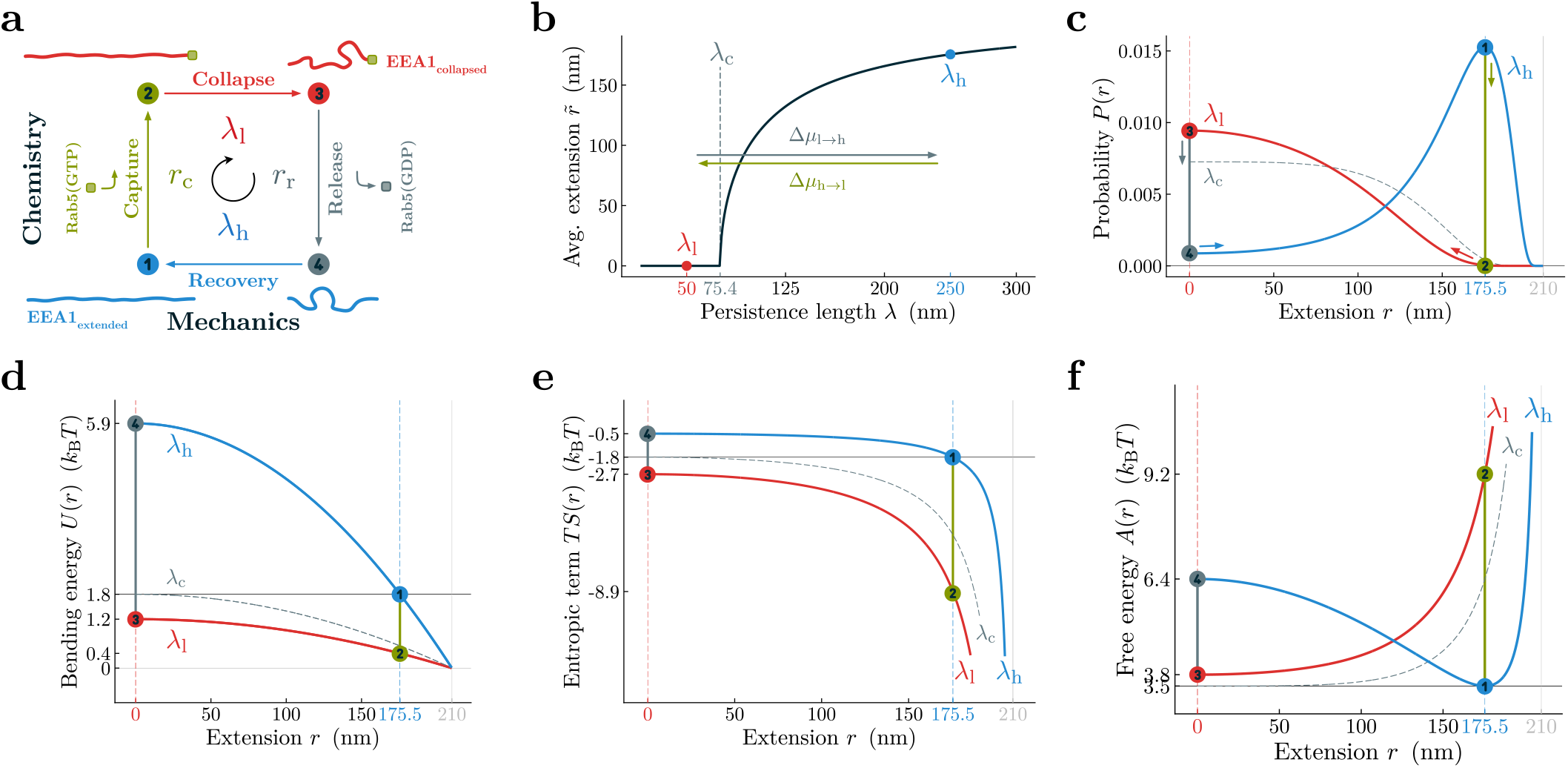
A two-state semi-flexible polymer model for the EEA1-Rab5 system. **(a)** The chemical potential differences provided by active or inactive Rab5 interactions (green - binding Rab5(GTP), grey - GTP hydrolysis, phosphate release, and unbinding of Rab5(GDP)) lead to changes in the effective persistence length of EEA1, thus coupling chemical transformations of the GTPase to tether mechanics. State transitions are indicated by respective numbers and colors: 1 – unbound and extended EEA1, 2 – extended EEA1-Rab5 complex, 3 – collapsed EEA1-Rab5 complex, and 4 – unbound and collapsed EEA1. Transition paths are: 1 → 2 – Capture (green), 2 → 3 – Collapse (red), 3 → 4 – Release (grey), 4 → 1 – Recovery (blue). The variables held constant during the transitions are indicated next to the respective transition paths. **(b)** With a fixed contour length *L*, only the persistence length *λ* determines the equilibrium behaviour, like the average extension. The size of the system determines a critical persistence length *λ*_*c*_ = 2*π*^−3/2^ *L* above which the equilibrium extension is greater than zero. The two states of unbound and bound EEA1 toggle around this critical value. **(c)** The probability density of the extension illustrates how the substantial shift between high and low persistence lengths (from *λ*_h_ = 250 nm (blue) to *λ*_l_ = 50 nm (red), *L* = 210 nm) affects the expected end-to-end tether extension in the constant length ensemble. Indicated states are color-coded as in **(a)**. Arrows designate the direction of the transition paths. For comparison, the probability density for the critical persistence length is also shown (grey, dashed curve). **(d-f)** The bending energy, *U* (*r, λ*) **(d)**, and the conformational entropy, *S*(*r, λ*) **(e)**, contribute to the free energy, *A*(*r, λ*) **(f)**, of the system. The various states and transitions of the molecular motor system are illustrated in each panel with the same color-coding as in **(a)**. Together, these panels indicate the asymmetry of the individual motor substeps due to the semiflexible nature of the tether: whereas the collapse is entropically driven, the recovery is dominated by the energetic contribution. Furthermore, at their respective equilibrium extensions, it costs more free energy to turn a stiff polymer into a more flexible one, than *vice versa*.

Our theoretical analysis is performed in a ‘fixed extension ensemble’ where the end-to-end distance of the EEA1 molecule is fixed on shorter timescales but is allowed to slowly vary as the molecule proceeds through the mechanochemical cycle. Key elements of the cyclic protocol involve four processes that comprise a closed path (Fig. 3**a**; the cycle is indicated by the lines connecting the points `1,2,3` and `4`): 1) Upon binding to Rab5(GTP), EEA1 reduces its bending rigidity *κ* and its persistence length decreases from *λ*_h_ to *λ*_l_ (upward green arrow). 2) The extended but soft semi-flexible polymer equilibrates mechanically, resulting in a reduction of its end-to-end distance (red arrow, collapse). 3) Triggered via a combination of chemical steps that include GTP hydrolysis by Rab5, phosphate release and the unbinding of Rab5(GDP) from EEA1, EEA1 undergoes a switch back from soft to stiff, *i*.*e*. its persistence length increases from *λ*_l_ to *λ*_h_ (upward grey arrow). 4) The now collapsed but rigid semi-flexible polymer equilibrates mechanically resulting in an extension of its end-to-end distance (blue arrow). Two segments of the path, *i*.*e*. `2`→`3` (collapse) and `4`→`1` (recovery), amount to a change of extension that can proceed against an external opposing force, and EEA1 can perform mechanical work in these segments. Transitions between the two flexibility states of EEA1, *i*.*e*. `1`→`2` and `3`→`4`, are mediated by the aforementioned chemical steps. The chemical potential differences Δ*µ*_*h*→*l*_ and Δ*µ*_*l*→*h*_ capture the free energy differences of the respective chemical transitions within the EEA1-Rab5 cycle. Taken together, EEA1-Rab5 is able to convert chemical energy to mechanical work during one passage of this simplified mechanochemical cycle.

We next discuss how EEA1-Rab5 can perform mechanical work in the fixed extension ensemble to which the protocol of slowly varying extension described above is applied. The EEA1 molecule is held in place between two attachment points that are fixed in space, *i*.*e*. with a fixed end-to-end distance, while the force that the molecule exerts on the attachment points fluctuates. The two attachment points exert equal but opposite external forces in order to keep their positions fixed in space. As the cyclic protocol proceeds, the molecule is subjected to a slow change in attachment point positions, leading to a slow change of the end-to-end distance of EEA1 concomitant with a change in the average force that the molecule exerts on the attachment points. We note that the switch of bending rigidity, *κ* upon Rab5 binding (`1`→`2`) and release (`3`→`4`) of Rab5 causes EEA1 to transition between two regimes: *λ* ≪ *L* when flexible and *λ* ≳ *L* in the more rigid state. This requires us to utilize the Blundell-Terentjew model for semiflexible polymers that is valid in both regimes (19).

We highlight a key aspect of this model that makes it suitable for both flexibility regimes. In an isothermal environment, for a polymer with a fixed contour length, *L*, the average equilibrium end-to-end distance, 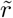 is determined by the effective persistence length, *λ*. The model identifies a critical persistence length, *λ*_*c*_ = 2*π*^−3/2^*L*, below which 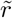 becomes zero (Fig. 3**b**) and the polymer behaviour is effectively Gaussian. This critical point therefore separates the regimes of high and low flexibility, as we have also identified for EEA1 in the FCS experiments (Figs. 1 and 2).

This transition therefore suggests a qualitative difference, *i*.*e*. a buckling transition between the two regimes for a semi-flexible polymer: one where the filament can withstand thermal fluctuations (*λ > λ*_c_) to maintain a finite extension, and one where it cannot and therefore has an equilibrium end-to-end distance of zero.

Experiments performed here and in previous work (7), indicate that EEA1 is indeed poised close to such a critical point. The unbound, extended state of EEA1 and the Rab5(GTP) bound, collapsed state indeed lie on either side of the critical persistence length *λ*_c_ ≈ 75 nm for the *L* ≈ 210 nm long molecule (Fig. 3**b**). Altogether, this suggests a mechanical state transition in EEA1 mediated by its chemical interaction with Rab5. For our subsequent analysis (and without any loss of generality) we take *λ*_h_ = 250 nm and *λ*_l_ = 50 nm, which are the consensus values from various experiments (dcFCCS here, and optical tweezer, rotary shadowing EM (7)) for the extended and collapsed states, respectively.

To compute the forces exerted by EEA1, we first consider the conformational free energy of a polymer in the absence of chemical transitions. We start from the analytical form for the polymer end-to-end distance distribution, *P* (*r, λ*). The Helmholtz conformational free energy of EEA1 is related as *A*(*r, λ*) = −k_B_Tlog P(r, *λ*, L), from which the forces exerted by the molecule are computed as ⟨*F*⟩ (*r, λ, L*) = −*∂*_*r*_*A*(*r, λ, L*). The free energy *A*(*r, λ*), and specifically the balance between the bending energy *U* (*r, λ*) and the conformational entropy *S*(*r, λ*) (Fig. 3**d,e**), reveal interesting aspects of the EEA1-Rab5 motor. Before discussing the relative contributions of the entropic and bending energy components, it can already be seen that the mechanical transitions are driven by the minimization of the polymer free energy (Fig. 3**f**). This minimization can be understood as follows: the persistence length switch along the path `1`→`2` is associated with a reduction in both the conformational entropy *S* and bending energy *U* of EEA1 (Fig. 3**d,e** and Supplementary Information Sec. III). The collapse of EEA1, *i*.*e*. `2`→`3` is therefore entropic in origin, and the mechanical force generation is akin to that of an entropic spring (25). In contrast, the recovery of EEA1 to the extended configuration along the path `4`→`1` is driven by the stored polymer bending energy, which results in a pushing force. In equilibrium, the unbound polymer has a finite extension that exactly balances the entropic and energetic contributions. As expected, all states of a semi-flexible polymer fall on a universal curve that asymptotically approaches the two extremes: a rigid rod (fully elastic) or a flexible freely jointed chain (Supplementary Information Sec. III).

From the forces, we estimate the mechanical work performed by the two-component motor, for which we consider two generic scenarios for the work cycle: (i) EEA1 is tethered to a vesicle cargo which it can drag within the dense cytoplasmic milieu and (ii) an *in vitro* situation in which EEA1 is tethered to a colloidal probe (spherical bead) and exerts a force against an optical tweezer that holds the colloid (7). We compute the average force at a given extension, ⟨*F*⟩ (*r, λ, L*) = − *∂*_*r*_*A*(*r, λ, L*) and this force-extension relationship (shown for the two persistence lengths associated with the extended and collapsed states of EEA1 in Fig. 4**a**) corresponds to having an external force *F*_ext_ = −⟨*F* (*r, λ*)⟩ that maintains the polymer end-to-end extension *r* fixed. In the fixed extension ensemble discussed above, the capture and release extensions are variable, and their magnitude, together with the persistence length corresponding to the two flexibility states of EEA1, determines the amount of work that is extracted. In the first scenario *i*.*e* the context of cellular vesicle trafficking, EEA1 is most likely to capture a vesicle at the average equilibrium extension of the unbound tether (*r*_capture_ ≈ 175.5 nm) and collapse until other molecular layers start to engage with the vesicle (*r*_release_ ≈ 75 nm) leading to docking and completion of membrane fusion. From here the tether recovers and could potentially push out a load following the blue curve to reach the unbound equilibrium extension, after which the cycle could start again with a new round of vesicle tethering. On the other hand, analogous to the *in vitro* situation of EEA1 dragging a tethered colloid against an optical tweezer, there is a time-dependent tension and drag force (26), during the entropic collapse, against which EEA1 performs work (Fig. 4**b**). The mechanical work computed from our model, quantitatively matches with the mechanical work distribution that was measured in optical tweezer single-molecule experiments (Fig. 4**c**). Based on the optical tweezer measurements and the different combinations of persistence lengths in the extended and collapsed states (Supplementary Information Sec. III), the maximum work that can be obtained by entropic collapse mechanism ranges between 6 and 20 k_B_T, depending on the relative differences in flexibility of EEA1 in the two states (detailed investigations for different values of the persistence length are shown in the Supplementary Information Sec. III).

**Fig. 4.**
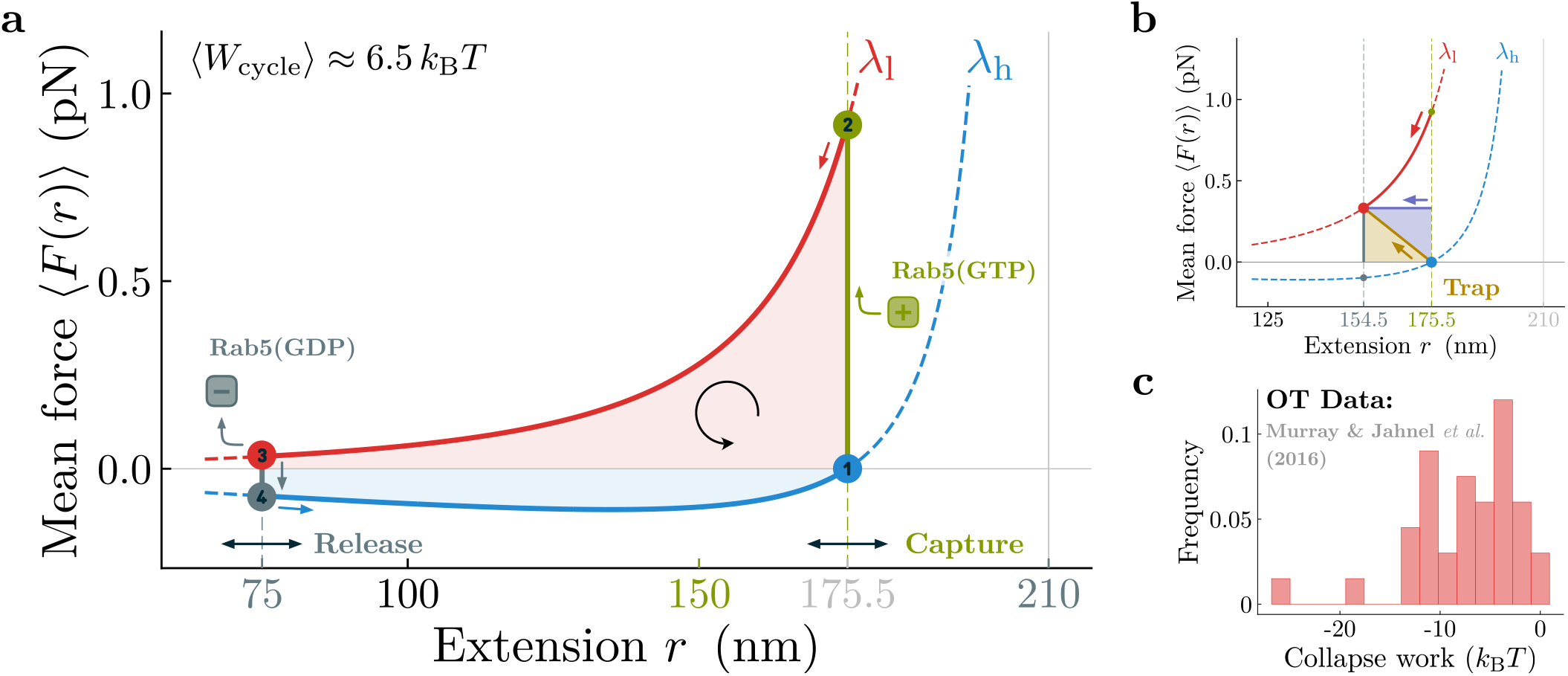
A switchable polymer engine can perform work against an external load. **(a)** With the switchable semi-flexible polymer as the working substance, cyclic motor-like processes can be run by corresponding protocols. One example is an idealized Stirling-type engine. Here, the four steps of the work cycle are as follows: 1 → 2 – isometric softening after active Rab5 capture without performing work, 2 → 3 – isothermal collapse (red) to perform work (red shaded area), 3 → 4 – isometric stiffening after release of inactive Rab5 (grey). 4 → 1 – isothermal extension (blue) to perform work during recovery (blue shaded area). This portion might be harder to extract due to the detachment of the GTPase. The indicated capture extension (green) corresponds to the equilibrium extension of the unbound tether, while the release extension (grey) is variable and has been chosen for illustrative purposes. **(b)** Coupling the motor to an external force, like an harmonically trapped microsphere (yellow) or a constant drag force (violet), mechanical work can be performed. **(c)** Relative occurrence of measured collapse work in single-molecule EEA1-Rab5 optical tweezer experiments from Ref. (7)

For this two-component system to operate as a self-sufficient GTP-fuelled motor, the increase in free energy from `1`→`2` and `3`→`4` must be accounted for by the chemical energy transitions arising from chemical potential differences and the associated chemical transitions due to Rab5(GTP) binding/unbinding, GTP hydrolysis and phosphate release, and the unbinding of Rab5(GDP). These chemical free energies, combined into the effective chemical potentials Δ*µ*_h→l_(*r* = *r*_capture_) and Δ*µ*_l→h_(*r* = *r*_release_), set upper bounds on the mechanical work that the EEA1-Rab5 motor can perform. It is important to note that for EEA1-Rab5 to proceed through the cycle without the aid of external factors the sum of the two chemical transitions Δ*µ*_total_ = Δ*µ*_h→l_(*r*_capture_) + Δ*µ*_l→h_(*r*_release_) cannot exceed the available chemical energy per cycle derived from GTP hydrolysis Δ*µ*_total_ < Δ*µ*_GTPhydrolysis_. The dissociation constant between Rab5(GTP) and the Zinc-finger domain of EEA1 is *K*_D_ = 2.4 µM (23), which translates into a free energy of 12.9 k_B_T for Rab5(GTP) binding. Several lines of evidence suggest that the binding between active Rab5 and the full length dimeric EEA1 is even stronger (27), supporting the assumption that the binding step could provide enough free energy to drive the flexibility transition (transition from 1 → 2 in Fig. 3**a**). Meanwhile, the hydrolysis of a single Rab5(GTP) can produce an energy equivalent to 20−25 k_B_T (28). Since the energy of GTP hydrolysis is larger than the energy obtained from binding, it is feasible that the sum of both chemical potentials (capture and release, even considering that according to the model release would require ≈ 3 k_B_T, see Fig.3**f**) are lower than the energy obtained from GTP hydrolysis. Altogether, while the polymer model allows us to compute the mechanical work performed by EEA1 during the entire cycle, the chemical energy obtained from binding, GTP hydrolysis, phosphate release and unbinding sets upper bounds for this work, which can be as high as 20−25 k_B_T (28). The energy released by either the GTP hydrolysis and or the Rab5(GTP) binding would be sufficient to account for the work performed during a single cycle of EEA1 extension and collapse for the persistence length values that we have measured in the various experiments (dcFCCS here and optical tweezers and rotary shadow EM earlier (7)). Conversely, this limits the allowable transitions in the bending rigidity, *κ*, of the extended and collapsed configurations of EEA1. Finally, since the EEA1–Rab5 system requires two chemical energy transitions (*i*.*e*. `1`→`2` and `3`→`4` in Fig. 3**a** and Fig. 4**a**, both free energy sources (Rab5(GTP) binding and GTP hydrolysis) can be utilized to complete a full cycle. Finally, the efficiency of this motor (Extended Data Fig. 9) is in the range of other molecular motors such as kinesins (14).

## Discussions

While the roles of small GTPases in cellular signal transduction have long been recognised, their function in mediating mechanical processes has thus far remained less explored. On the other hand, many ATPases like myosin and kinesin have been recognised as force-generating soft machines for several decades, although they are rarely seen as signalling molecules. Together with our earlier results (7), this work demonstrates that the coiled-coil protein EEA1 and the small GTPase signalling molecule Rab5 work together as a two-component molecular motor system that can transfer the chemical energy of GTP hydrolysis into mechanical work, while simultaneously toggling the state of the GTPase during the process. Altogether, this sets this GTPase driven mechanism apart from other ATP-driven motors (14).

Our results have important implications for membrane tethering and fusion well beyond the specific case of EEA1 and Rab5. Several long coiled coil tether proteins that function at distinct stages of the exocytic and endocytic pathways are effectors of small GTPases (29–32) — these long molecules have traditionally been described as *spacers, rulers* (33), and *rods* (34), suggesting static and fixed distances between the ends. Accordingly, the observation that the physical length of a coiled-coil domain is much more conserved than its sequence (35) has been interpreted along the lines that a static distance between the two ends is functionally important. Yet, contrary to the widely accepted notions of coiled-coils as static and fixed rulers (33), our work shows that EEA1, a prominent example of a long coiled-coil tethering molecule, is poised at a critical point close to bistable flexibility states. The switch between these states is mediated by a series of chemical steps that induce large-scale movements on the order of ≈ 100. The mechanism described here could be shared by several other protein tethers. The Golgi coiled-coil tether such as GCC185 exhibits a great degree of flexibility due to unwinding of the coiled coil central region (8). This supports the idea that long coiled coil tethers are “breathing” molecules, *i*.*e*. metastable mechanical structures. Although in that case the effect of binding to Rab GTPases has not been explored yet, it is possible that it may control the dominant conformation of the tether, as shown for EEA1. This would make the tether-small GTPase coupling a widespread paradigm not only for membrane recognition but also to convert the energy stored in GTP into work in the mechanochemical pathways to membrane fusion. If a work cycle were a general requirement of the membrane tethering-to-fusion process, the multi-protein tethering complexes such as Exocyst and HOPS could use mechanical energy as part of their function in a similar fashion to the long coiled coil tethers. However, in contrast to flexibility transitions, one could expect this to be an enthalpy-driven transition i.e. a lever arm like action at a single location as opposed to an overall “softening” of the protein.

Finally, the model we identified to harness mechanical work from this cyclically acting polymer motor (Fig. 4**a**) is reminiscent of a classical Stirling engine. While classical engines use gas as the working substrate and exploit temperature gradients to generate mechanical work via expansion/compression of the gas, the two-component polymer motor that we describe here uses the polymer itself as working substrate with polymer flexibility changes resulting in force generation. Consequently, this kind of mechanism suggests new avenues for the design of synthetic polymer engines (36).

## ACKNOWLEDGEMENTS

We thank Madan Rao and Pramod Pullarkat for discussions. We acknowledge support from the Department of Atomic Energy, Government of India, under projects RTI4001 and RTI4006, the Simons Foundation (Grant No. 287975 to ST) and the Max Planck Society through a Max-Planck-Partner-Group at NCBS-TIFR (ST). MJ and SWG were Supported by the Deutsche Forschungsgemeinschaft (DFG, German Research Foundation) under Germany’s Excellence Strategy – EXC-2068 – 390729961. JAS and JL were financially supported by the Human Frontiers in Science Program (HFSP), grant number RGP0019/2020. This study was financially supported by the Max Planck Society. We thank the following Services and Facilities of the MPI-CBG for their support: PEPC and Light Microscopy. We also thank the Central Imaging and Flow Cytometry Facility (CIFF) at the NCBS.

## Author Contributions

MJ conceived the two component molecular motor idea. ST, AS, JAS, MZ and JL conceived and designed the FCS experiments. JAS and JL did the biochemistry and sample preparation. AS and JAS performed the FCS experiments. AS analyzed the FCS data with guidance from ST. MJ analyzed the optical tweezers data. JAS, JL and MZ contributed knowledge of the GTPase cycle to the development of the physics model. MJ developed the polymer physics model, with help from ST and SWG. All authors interpreted the data and contributed to the writing of the paper. ST, MZ and SWG provided overall supervision for the project.

## Materials and Methods

### Cloning, expression and purification of proteins

EEA1 was sub-cloned into the pOEM1-based vector pOCC151 (PEPC; MPI-CBG), which includes an N-terminal GST tag followed by a HRV-3C cleavable site between the tag and the inserted gene. On the EEA1 C-terminus the amino-acid sequence “GGGSGGGGSGGGGSGGGGSLPETGGGG” was added. Rab5a was sub-cloned into the pET11-based vector pOCC9 (PEPC; MPI-CBG), which includes an N-terminal hexahistidine tag followed by a HRV-3C cleavable site between the tag and the inserted gene. The plasmid pET30b-7M SrtA, containing a C-terminal hexahistidine tag, was a gift from Hidde Ploegh (Addgene plasmid 51141).

Rab5 and SrtA7m were expressed in E.coli-BL21(DE3) and protein purification was performed in standard buffer (20 mM Tris pH7.4, 150 mM NaCl, 5mM MgCl_2_, 0.5 mM TCEP) as specified in Supplementary Information. For GTP loading, freshly purified Rab5 was supplemented with 10 mM EDTA and 10-fold excess GTP. The mixture was incubated for 15 minutes at 4°C before addition of 10 mM MgCl_2_, and subsequently ran over a desalting column equilibrated in standard buffer. EEA1 was expressed in SF9 cells growing in ESF921 media (Expression Systems) as specified in Supplementary Information Sec. I.

### Dual color labelling of EEA1

The SrtA recognition site (LPETG) was added to the EEA1 C-terminus during sub-cloning. The SrtA-based reaction was performed in buffer containing 20 mM Tris pH 7.4, 150 mM NaCl and 0.5 mM TCEP. A mixture of 1 *µ*M EEA1, 30 *µ*M GGGaWC-A488, 30 *µ*M GGGaWC-A647 and 1.5 *µ*M SrtA7m was incubated for 1 hr at room temperature on a rotator wheel. The obtained EEA1-fluorophore conjugate was purified by size exclusion chromatography and the purity was evaluated by SDS-PAGE, laser scanning imaging (Typhoon FLA9500; GE Healthcare; 473nm and 635nm wavelength) and coomassie staining. GGGaWC-A488 and GGGaWC-A647 were produced by the Biomolecular Synthesis facility at BCUBE (Dresden). For more detailed information refer to Supplementary Information Sec. I.B.

### dcFCCS setup and equipment

dcFCCS experiments were performed on the Confocor3 setup (Carl Zeiss LSM 780 NLO) operated in photon counting mode[objective: C-Apochromat 40× 1.2 NA W Corr M27; excitation sources: 488*nm* (Ar-ion laser: 4.52*µW*) and 633*nm* (HeNe laser: 6.23*µW*); emission range for detectors: 499 − 552*nm* and 641 − 694*nm*; pinhole diameter: 58*µm*; temperature: 23 ± 0.5°*C*]. 10*nM* dsDNA (50*µl*), dual labelled with A488 and A647 at opposite ends, was used to maximize confocal volume overlap by adjusting the objective collar for maximum photon counts per molecule. Confocal volume alignments in the XY- and Z-planes were visualized by confocal microscopy of fluorescent membrane coated beads at their equatorial plane and by a Z-scan of supported lipid bilayers respectively, where membranes were doped with fluorescent membrane probes DIO & DID (Supplementary Information Sec. II).

### Dynamics of a single polymer end

The correlation *G*(*τ*) of the fluctuating labelled end is related to the MSD(⟨*r*^2^(*τ*)⟩) of the end of the polymer

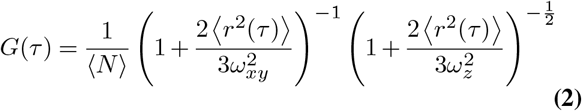

⟨*N*⟩, *ω*_*xy*_ and *ω*_*z*_ correspond to the average number of molecules in the confocal volume, the lateral and the axial width of the confocal volume. Normalized correlation curves were fitted to Eq. 2 and the roots (⟨*r*^2^(*τ*)⟩) were obtained by Broyden’s good method using a custom written computer code. The local exponent *α*(*τ*) was computed by finding the local slope for a moving window of ⟨*r*^2^(*τ*)⟩ corresponding to a decade of lag-time *τ* using expression:

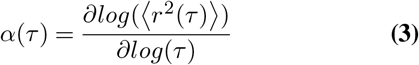

By fitting (⟨*r*^2^(*τ*)⟩) to Eq. 1 for *n* ≤ 2, the diffusion coefficient *D*, time-scales pre-factor Δ_*n*_ and time-scales (*τ*_*n*_) were obtained. The persistence length *λ* was calculated by fitting to expanded form (Eq. 4) of Eq. 1, written as a function of radius of cross-sectional area (*a*), contour length (*L*), and persistence length(*λ*) of the polymer

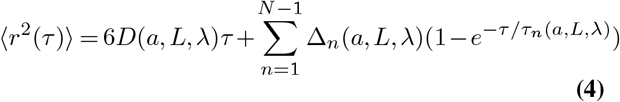

The parameters, *a*(= 1*nm*) and *L*(= 220*nm*) were obtained from the crystal structure of EEA1 coiled-coiled region (*PDB* : 1*JOC*) and rotary shadow electron microscopy from Murray *et al*.respectively. Using *a* and *L* as fixed parameters in Eq. 4, results in the free parameter *λ*.

### Bootstrapping

The recovery from collapsed to extended state results in intermediate populations which were probed by performing bootstrapping on the dcFCCS curves. Bootstrapping was performed on 180 curves. Mean correlation (*G*(*τ*)^*avg*^) curve was obtained from 20 curves selected at random. 10^5^ such *G*(*τ*)^*avg*^ curves were sampled with replacement to calculate the distribution of *λ*. For intermediate stages of recovery, multi-modal distributions were obtained for the two cycles with peaks corresponding to collapsed and extended states (more details can be found in Supplementary Information Sec. II.E4).

### Experiments to probe EEA1 flexibility in different states

Experiments were carried out in a 384-well glass bottom plate with 175±15*µ*m glass thickness sealed with aluminium foil to prevent drying. 50*µ*l of EEA1 (100*nM*) were added to a well and the confocal beams were focused 20*µ*m above the glass surface to perform dcFCCS. 240 correlation curves of 30 seconds each were recorded for 2 hours. The first 30 minutes recordings were removed to let the system equilibrate. Two rounds of 2*µ*M Rab5(GTP) were added to the solution and dcFCCS was evaluated for 2.5 hours after each addition. The data was further analyzed as described in Supplementary Information Sec. II.

### Collapse work estimation from single molecule optical tweezer experiments

To estimate the order of magnitude of mechanical work performed during Rab5-driven collapse of EEA1, we re-analyzed single molecule optical tweezer data from our earlier study. In brief, for these experiments, glass beads with supported lipid bilayers carrying purified EEA1 or active Rab5(GTP) were trapped in a dual-trap optical tweezer. The traps were successively brought closer together until there was a connection between an EEA1 on one microsphere and Rab5(GTP) on the other one. We observed transient interactions which pulled both beads together and resulted in sub-pN forces (see Figure 3 of Ref. (7)). Using the known trap stiffness and displacements for each experiment, here we calculated the mechanical work for each event for the case of pairing EEA1 with Rab5(GTP) covered beads (N = 38). We interpret the measured work in the optical tweezer as corresponding to the transition 1 → 2 → 3 in Fig. 4**a** which amounts to mechanical work performed during entropic collapse. The histogram of this collapse work is shown in Fig. 4**c**.

