## Supplementary Text and Figures for "GTPase-dependent cyclic flexibility transitions drive the two-component EEA1-Rab5 molecular motor"

### I. PROTEIN PREPARATION AND ENZYME BASED FLUORESCENT LABELLING

#### A. Expression and purification of proteins

Rab5 and SrtA7m were transformed in E.coli-BL21(DE3) and protein expression was induced with 50mM Lactose. After 16 hours incubation at 30°C the culture was spun for 20 minutes at 4000g using a JLA 8.1000 rotor. The pellet was resuspended in standard buffer (20mM Tris pH7.4, 150mM NaCl, 5mM $MgCl_2$, 0.5mM TCEP) and supplemented with DNase I (Sigma) and protease inhibitor cocktail (chymostatin 6 $\mu$g/ml, leupeptin 0.5 $\mu$g/ml, antipain-HCl 10 $\mu$g/ml, aprotinin 2 $\mu$g/ml, pepstatin 0.7 $\mu$g/ml, APMSF 10 $\mu$g/ml) before being flash frozen and stored at -80°C.

EEA1 was expressed using a Baculovirus Expression System. SF9 cells, cultured in ESF921 media (Expression Systems), were co-transfected with the DNA construct containing the EEA1 gene and linearized viral genome. After selecting for high infectivity, P1 and P2 virus were generated as described on the manufacturers protocol. SF9 cells at 1 million $cells/ml$ were infected with P2 virus at 1% vol/vol and shaken at 27 °C for 48 hours. Cells were harvested by spinning at 500g for 10 minutes. As with bacteria-expressed proteins, the pellet was re-suspended in standard buffer enriched with DNase I and protease inhibitor cocktail before being flash frozen and stored at -80°C. The pellet was thawed on ice and all subsequent steps were performed on ice or at 4°C. E.coli cells were lysed via sonication and SF9 cells via Dounce homogenizor. Lysates were ultracentrifuged for 1 hour at 142000g with Ti45 rotor (Beckman Coulter). For histidine-tagged protein capture, the supernatant was incubated for 1 hour with Ni-NTA Agarose resin (Qiagen), and for GST with GS-4B resin (GE Healthcare) for 2 hours. Following several washes with standard buffer, histidine-tagged proteins (Rab5 and SrtA7m) were eluted with 200mM Imidazole, and GST-tagged EEA1 was cleaved from the resin by overnight incubation with HRV-3C protease. The proteins were further purified by size exclusion chromatography on Superdex200 (Rab5 and SrtA7m) or Superose6 (EEA1). Purity was evaluated by SDS-PAGE followed by comassie staining, and the concentration measured by BCA assay (ThermoScientific). Finally, the protein was aliquoted, flash frozen in liquid nitrogen and stored at -80°C. The fluorescent labelling of EEA1 was performed following purification as described in I B.

##### B. Enzyme-based labelling of EEA1 with Alexa488 and Alexa647

S.aureus Sortase A (SrtA) is a transpeptidase that recognizes the peptide sequences (LPXTG; where X is any amino acid) and (GGG) and forms a covalent bond between them [1]. We added the SrtA recognition-site (LPETG) on EEA1 C-terminal while the (GGG) was linked to either Alexa488 or Alexa647. The heptamutant version of the SortaseA enzyme 'SrtA-7m' was chosen because it is $Ca^{2+}$ independent and has an increased efficiency compared to the wild type [2].

The EEA1 modification includes a linker, the SrtA recognition-site and three extra aminoacids. The linker and extra C-terminal aminoacids were included to improve the accessibility and affinity of the enzyme to its recognition site [3][4]. Besides, the relatively long linker (19 amino-acids) also helps avoid any possible cross-talk or FRET pairing between the fluorophores.

The conjugation between the GGG peptide and the alexa dyes was performed by the Biomolecular Synthesis facility at BCUBE (Dresden). The compounds generated were "GGGaWC-(Alexa488)" and "GGGaWC-(Alexa647)". "a" is a beta-alanine acting as a linker. The SrtA-based reaction was performed in buffer containing 150$mM$ NaCl, 20mM Tris pH 7.4 and 0.5$mM$ TCEP. A mixture of 1$\mu$M EEA1, 30$\mu$M GGGaWC-A488, 30$\mu$M GGGaWC-A647 and 1.5$\mu$M SrtA-7m was incubated for 1 hour at room temperature in a rotator wheel. The labelling of each monomer is independent, resulting in EEA1 dimers labelled with two Alexa488, two Alexa647 or the combination of one Alexa488 and one Alexa647 (Extended data Fig. 1a). The EEA1-fluorophore conjugate was purified by size exclusion chromatography and the purity evaluated by SDS-PAGE, laser scanning imaging (Typhoon FLA9500; GE Healthcare; 473nm and 635nm wavelength) and coomassie staining (Extended data Fig. 1b).

#### II. DUAL COLOR FLUORESCENCE CROSS-CORRELATION SPECTROSCOPY

##### A. Oligo annealing

dsDNA labelled with Alexa488 and Alexa647 at opposing ends was used to optimize the objective collar setting for maximizing confocal volume overlap and further calculating the

overlapped confocal volume. More details in II D. To generate dsDNA labelled with two fluorophores was followed the same strategy and oligonucleotide sequence as in [5]. Two complementary DNA oligonucleotides were synthesized, labelled at their 5' end with either Alexa647 or Alexa488 and HPLC purified by Eurofins. The sequences are:
Alexa647-5'-ATGGCTAATGACCGAGAATAGGGATCCGAAT TCAATATTGGTACCTA CGGGCTTTGCGCTCGTATC-3' and Alexa488-5'-GATACGAGCGCAAAGCCCGTAGG TACCAATATTGAATTCGGATCCCTATTCTCGGTCATTAGCCAT-3'.

Annealing of the complementary strands was performed at 10$\mu$M oligonucleotide concentration in a buffer containing 100$mM$ KOAc, 25$mM$ Tris-acetate (pH 7.6), 10$mM$ MgOAc, 0.5$mM$ $\beta$-mercaptoethanol and 10$\mu g/ml$ BSA. The solution was heated in a thermocycler at 95°C for 2 minutes and slowly cooled down to 23°C using a temperature gradient of 1.2°C/min.

##### B. Membrane coated beads and supported lipid bilayers preparation

Membrane-coated-beads (MCBs) and supported-lipid-bilayers (SLBs) were used to evaluate the confocal volume overlap between 488nm and 633nm channels on the xy-plane (MCBs) and z-axis (SLBs). More details in II D.

Lipids were stored at -20°C in chloroform. For liposome formation, 700 nanomols of lipid were mixed at a molar ratio of "99.8% DOPC (Avanti), 0.1% DID (ThermoFischer), 0.1% DIO (ThermoFischer)" and the solvent evaporated under nitrogen flow followed by overnight drying under vacuum. Lipids were rehydrated in 700$\mu$l SLB buffer (20$mM$ Tris pH 7.4, 150$mM$ NaCl) at 37°C and vortexed to achive a 1$mM$ stock. Small uni-lamellar vesicles (SUVs) were formed by 10 freeze-thaw cycles including submersion in liquid nitrogen, 5 minutes in a water bath at 37°C, and stringent vortexing. SUVs were sonicated with a tip sonifier (microtip 450D, Branson) at an amplitude of 34% for 1 minute with 1.5 seconds pulses ON/OFF.

To form membrane coated beads (MCBs), 10$\mu$M silica beads (140244-10, Corpuscular) at 10 $million/ml$ were incubated with 400$\mu$M liposomes and 800$mM$ NaCl for 30 minutes at room temperature on a rotator wheel. Obtained MCBs were sequentially washed with 20$mM$ Tris at pH7.4 and standard buffer by 1 minute centrifugations at 380$g$ in a tabletop centrifuge. Formation of MCBs was optimized to have excess membrane reservoir. Upon sedimentation, that reservoir spilled onto the glass surface, effectively forming a supported lipid bilayer (SLB). With this strategy we could perform alignment in the xy-plane and z-axis using the same chamber, thus minimising technical difficulties. MCBs and SLBs were imaged within 1 hour since formation.

##### C. Experimental Setup and equipment

FCS and dcFCCS experiments were performed on Carl Zeiss LSM 780 NLO inverted microscope with ConfoCor3 setup using "C-Apochromat 40× /1.2 W Corr M27". This objective was chosen to avoid chromatic and spherical aberrations and to achieve maximum confocal volume overlap for $488nm$(Argon) and $633nm$(HeNe) lasers. Selected detectors have a range of $499 - 552nm$ and $641 - 694nm$ respectively and are operated in photon counting mode. The time-intensity traces and the auto(cross)-correlations were recorded and obtained from Zen blue 10 software.

##### D. Optical path and confocal volume alignment

dsDNA (dual labelled with Alexa488 and Alexa647) was used for pinhole alignment in the optical path. $50\mu l$ dsDNA ($10nM$) was loaded to a well of a 384-well glass bottom plate. The fluorophores were first excited by $488nm$ laser, and the emission path aligned by maximizing the intensity on the detector ($499 - 552nm$ range: green channel) by moving the pinhole in the xy-plane. Similarly the process was repeated for the $647nm$ laser ($641 - 694nm$ detector range: red channel), resulting in aligned optical path.

Once the optical path is aligned, we further increase the confocal volume overlap by finding the appropriate objective collar setting. For that, the dual-labelled dsDNA was excited simultaneously with $488nm$ and $633nm$ lasers, and the photons emitted by both fluorophores (Alexa488 & Alexa647) collected in their respective detectors. The emitted photons were recorded as time-intensity traces for each channel. Time-intensity traces of each individual channel were auto-correlated for different lag-times to obtain auto-correlation curves. The time-intensity traces of green and red channels were cross-correlated for different lag-times to obtain cross-correlations curves. The objective collar position was optimized by maximizing the cross-correlation (G(0)) [6, 7].

The overlap in volumes was visualized by imaging 2-colour-labelled MCBs at their equatorial plane (alignment at the xy-plane, Extended data Fig. 2a-b), and by Z-scan of supported lipid bilayers (alignment at the z-axis, Extended data Fig. 2c-d). The position difference for the maximum intensities of the green and red channels is $20nm$ in the xy-plane and $283nm$ in the z-axis.

#### E. Data Analysis: Theory & curve fitting

##### 1. *From end monomer fluctuations to correlation curves*

End-monomer dynamics of the labelled end of a polymer was used in the past for studying the biophysical properties of biopolymers such as DNA and RNA. FCS is employed to study the dynamics of the fluorescently labelled end of the polymer. The correlation $G(\tau)$ of the fluctuating labelled end is directly related to the mean square deviation ($\langle\langle r^2(\tau)\rangle\rangle$) of the end of the polymer.

$$G(\tau) = \frac{1}{\langle N\rangle}\left(1 + \frac{2\,\langle r^2(\tau)\rangle}{3\omega_{xy}^2}\right)^{-1}\left(1 + \frac{2\,\langle r^2(\tau)\rangle}{3\omega_z^2}\right)^{\frac{-1}{2}} \tag{1}$$

$G(\tau)$ is the correlation at the lag time $\tau$. The mean square deviation (MSD) at that lag time is represented by $\langle r^2(\tau)\rangle$. $\langle N\rangle$, $\omega_{xy}$ and $\omega_z$ correspond to the number of molecules in the confocal volume, the lateral and the axial radius of the confocal volume respectively. The obtained correlation curves are used to calculate the mean square deviation (MSD) as a function of lag time using expression 1. To calculate the $\langle r^2(\tau)\rangle$, the correlation plots are normalized with $G(0)$, which effectively results in $N = 1$. $G(0)$ is calculated by taking the mean of $G(\tau)$ for the first 10 lag-time values, and with this, normalized FCS curves are obtained. MSD is calculated using the values from normalized correlation curves using equation 1 and obtaining the roots by Broyden's good method [8, 9] with the help of a custom written python code. The values are further plotted to find the change in the local slope $\alpha(\tau)$ of $\langle r^2(\tau)\rangle$ corresponding to the cross-over between internal motion and centre of mass motion.

$$\alpha(\tau) = \frac{\partial log(\langle r^2(\tau)\rangle)}{\partial log(\tau)} \tag{2}$$

The numerical calculation of the local exponent $\alpha(\tau)$ is performed by taking data points of $\langle r^2(\tau)\rangle$ corresponding to 1 order of magnitude of lag-time ($i.e$ 88 data points in current setup). The minima of $\alpha(\tau)$ correspsonds to the crossover from the internal modes to translational diffusion of the polymer.

##### 2. *Determining confocal volume for FCS and dcFCCS*

The confocal volumes are determined by using dsDNA labelled with two distinct fluorophores attached to opposite ends to avoid any FRET pairing. FCS and dcFCCS are performed on the same dsDNA sample. In FCS is evaluated the auto-correlation of a single channel, whereas in dcFCCS is assessed the cross-correlation of the two channels. An average correlation plot, obtained from 30 measurements of $30sec$ each, is used for the analysis in FCS and dcFCCS. The average correlation plot for FCS (dsDNA_A488) and dcFCCS (dsDNA_A488_A647) is shown in Extended data Fig. 3a. The correlation appears to decay slower for dsDNA_A488_A647 than for dsDNA_A488, a shift attributed to artefacts arising from non-perfect confocal volumes overlap in the dcFCCS [6, 7, 10]. The confocal beam width ($\omega_{xy}$) for FCS is calculated based on the literature reported diffusivity ($414\mu m^2/sec$) of Alexa488 at 25°C [11]. The mean time needed for A488 to diffuse out of the confocal volume, obtained from 10 FCS measurements, is $34.75 \pm 3.7\mu s$. The width of the confocal volume is calculated from the measured diffusion time ($\tau_D$) and the Alexa488's diffusivity using expression 3.

$$w_{xy}^2 = 4D\tau_D \tag{3}$$

Extended data Fig. 3a,b and c, are plotted considering the same confocal width for FCS and dcFCCS, which is not the case in practice, to visualize the artefacts of non-perfect overlap of confocal volume in dcFCCS. The $MSD(\tau)$ and $\alpha(\tau)$ were calculated as described in the previous section and are shown in Extended data Fig. 3b and c. In Extended data Fig. 3c the inflection points for the local slope are different between FCS and dcFCCS, which is expected because of the difference in confocal spot size. Nonetheless, the local slope at the inflection point is found to be similar $i.e$ 0.87 and 0.88 for FCS and dcFCCS respectively. To obtain the confocal volume of dcFCCS we make use of the maximum entropy fit for FCS and dcFCCS curves [8, 12]. The maximum entropy method predicts the likelihood of the timescale from the FCS data in a model independent fit for time-scale distribution of dsDNA dynamics. The application of maximum entropy fit results in a two-peak probability distribution for the time scales of both FCS and dcFCCS as shown in Extended data Fig. 3d. The two peaks for the distribution of time-scales are $102\mu s$ and $489\mu s$ for FCS, and $230\mu s$ and $980\mu s$ for dcFCCS. For a given polymer, the faster time-scales are attributed to the internal modes, whereas the slower time-scales are associated with translational diffusion. The ratios of the respective characteristic time of the peaks between FCS and dcFCCS are $1:4.794$ for the faster mode and $1:4.26$ for the diffusion. The confocal width of dcFCCS ($w_{xy}^{dcFCCS}$) is obtained by multiplying the confocal width in FCS ($w_{xy}^{FCS}$) with the ratio of diffusion time of dcFCCS ($\tau_D^{dcFCCS}$) to FCS ($\tau_D^{FCS}$).

$$w_{xy}^{dcFCCS} = \frac{\tau_D^{dcFCCS}}{\tau_D^{FCS}} w_{xy}^{FCS} \tag{4}$$

The calculated confocal width is $234.9nm$ for FCS and $333.5nm$ for dcFCCS. The structure parameters ($w_{xy}/w_z$), obtained from the fit of averaged dsDNA curves for FCS and dcFCCS, are 6.22 and 6.5 respectively.

##### *3. Persistence length ($\lambda$) calculation*

Calculation of $MSD(\tau)$ for the fluctuating fluorescently labelled end is described in the previous section. The $MSD(\tau)$ comprises of the time-scales associated with the internal modes and diffusion as referenced in equation 5

$$\langle r^2(\tau)\rangle = 6D\tau + \sum_{n=1}^{N-1} \Delta_n (1 - e^{-\tau/\tau_n}) \tag{5}$$

$D$ is the diffusion coefficient, $\Delta_n$ is the prefactor corresponding to the time-scale $\tau_n$. Equation 5 shows the relation between the time-scales of internal modes and the diffusivity. However the parameters $D$, $\Delta_n$ and $\tau_n$ are dependent on the physical properties of polymer, such as radius of cross-sectional area ($a$), contour length ($L$) and persistence length ($\lambda$) of the polymer. Hinczweski *et al.* make use of dynamic mean field theory to further express equation 5 in terms of the mentioned parameters $a$, $L$ and $\lambda$ [13].

$$\langle r^2(\tau)\rangle = 6D(a,L,\lambda)\tau + \sum_{n=1}^{N-1} \Delta_n(a,L,\lambda)(1 - e^{-\tau/\tau_n(a,L,\lambda)}) \tag{6}$$

$$D(a,L,\lambda) = k_B T \Theta_0 \Psi_0^2(L/2) \tag{7}$$

$$\Delta_n(a,L,\lambda) = 6k_B T \tau_n(a,L,\lambda) \Theta_n \Psi_n^2(L/2) \tag{8}$$

$$\tau_n(a, L, \lambda) \approx \Lambda_n^{-1} H_{nn}^{-1} - H_{nn}^{-2} \Lambda_n^{-1} \sum_{n \neq m} \frac{H_{nm}^2 \Lambda_m}{H_{nn}\Lambda_n - H_{mm}\Lambda_m} \tag{9}$$

$\Theta_n$ and $\Psi_n(S)$ are the diagonal elements of the interaction matrix $H_{nn}$ with eiganvalues $\Lambda_n$ and orthogonal functions defined by Hinczweski *et al.* [13].

Further, $\Theta_n$ is approximated to:

$$\Theta_n(a, L, \lambda) \approx H_{nn} + 2 \sum_{n \neq m} \frac{H_{nm}^2 \Lambda_m}{H_{nn}\Lambda_n - H_{mm}\Lambda_m} \tag{10}$$

and $\Psi_n(L/2)$ can be approximated to:

$$\Psi_n(L/2) \approx \psi_n(L/2) + \sqrt{\frac{1}{L}\frac{H_{0n}}{H_{nn}}} + \sum_{n \neq m} \psi_m(L/2) \frac{H_{nm}\Lambda_m}{H_{nn}\Lambda_n - H_{mm}\Lambda_m} \tag{11}$$

where $\psi_n$ represents the normal modes [13].

The variables defined in Eq. 6 are depenedent on $a$, $L$ and $\lambda$. Hence, we further express the interaction matrix $H_{nn}$ in terms of $a$, $L$ and $\lambda$. $H_{nn}$ can be simplified and approximated well within two regimes: $n, m \ll L/\lambda$ and $n, m \gg L/\lambda$.

For regime $n, m \ll L/\lambda$, the $H_{nn}$ is expressed as:

$$H_{nm} \approx 2a\mu_0\delta_{nm} + a\mu_0(I_{nm}^1 + I_{nm}^2) \tag{12}$$

The stokes mobility ($\mu_0$) of a single sphere of radius $a$ and viscosity $\eta$ is $\mu_0 = 1/(6\pi\eta a)$. Where,

$$I_{nm}^{(1)} \approx \begin{cases} \frac{4\sqrt{6}}{L\sqrt{\pi}} \left[2ae^{-3/2} - \lambda e^{-6a^2/\lambda^2} + a\sqrt{6\pi} erf(\sqrt{3/2}) - a\sqrt{6\pi} erf(\sqrt{6}a/\lambda)\right] \; n \neq m \\ \sqrt{\frac{6}{L}} \left[E_1(6a^2/\lambda^2) - E_1(3/2)\right] \; n = m \end{cases} \tag{13}$$

The exponential integral function ($E_v(z)$) is defined as $E_v(z) \equiv \int_1^\infty dt \; e^{-zt}/t^v$

$$I_{nm}^{(2)} \approx \begin{cases} -\frac{L}{\lambda} \frac{2\sqrt{6}}{(n+m)(\sqrt{n}+\sqrt{m})\pi^{3/2}} \; n \neq m \\ \sqrt{\frac{6L}{\pi\lambda}} \left(\frac{1}{n^{1/2}} - \frac{1}{2\pi n^{3/2}}\right) - 2\sqrt{\frac{12}{\pi}} \; n = m \end{cases} \tag{14}$$

$$\psi_n(s) \approx \begin{cases} (-1)^{(n-1)/2} \sqrt{\frac{2}{L}} sin(\frac{\pi n s}{L}) \; n \; odd \\ (-1)^{n/2} \sqrt{\frac{2}{L}} cos(\frac{\pi n s}{L}) \; n \; even \end{cases} \tag{15}$$

$$\Lambda_n \approx \frac{3k_BT\pi^2n^2}{2\lambda L^2} \tag{16}$$

$$\sum_{n\neq m} \frac{H_{nm}^2\Lambda_m}{H_{nn}\Lambda_n - H_{mm}\Lambda_m} \approx \sqrt{\frac{L}{\lambda}}\left(\frac{-108\pi+27\pi^2}{18\sqrt{6}\pi^{7/2}n^{3/2}} + \frac{-126+72\pi-16\sqrt{3}\pi}{18\sqrt{6}\pi^{7/2}n^{5/2}}\right) \tag{17}$$

$$\sum_{n\neq m} \psi_m(L/2)\frac{H_{nm}\Lambda_m}{H_{nn}\Lambda_n - H_{mm}\Lambda_m} \approx \sqrt{\frac{2}{L}}\left[\frac{1}{18}(-9+4\sqrt{3}) + \frac{-9+(9+\sqrt{3}\pi)}{27\pi^2 n} - \frac{72-45\pi+4\sqrt{3}\pi}{216\pi^3n^2}\right] \tag{18}$$

For regime $n, m \gg L/\lambda$

$$H_{nm} \approx 2a\mu_0\delta_{nm} + a\mu_0 I_{nm}^{(1)} \tag{19}$$

Where,

$$I_{nm}^{(1)} \approx \begin{cases} (m^2+n^2)\left(log\left(\frac{16a^2mn\pi^2}{L^2}\right)+2\gamma\right) \\ \frac{2\sqrt{6}}{\pi^{3/2}}\frac{(m+n-1)(m+n)^2\pi}{(m+n-1)(m+n)(m^2+n^2)} \; n\neq m \\ -\frac{\sqrt{6}}{n\pi^{3/2}}\left[3+n\pi\{3\gamma+E_1\left(\frac{3}{2}\right)+2log\left(\frac{\sqrt{6}a\pi n}{L}\right)\}\right] \; n=m \end{cases} \tag{20}$$

Where $\gamma \approx 0.5772$ is Euler's constant

$$\psi_n(s) \approx \begin{cases} (-1)^{(n-1)/2}\sqrt{\frac{2}{L}}sin(\frac{\pi(2n-1)s}{2L}) + \sqrt{\frac{1}{L}}\frac{sinh(\pi(2n-1)s/2L)}{cosh(\pi(2n-1)/4} \; n\ odd \\ (-1)^{n/2}\sqrt{\frac{2}{L}}cos(\frac{\pi(2n-1)s}{2L}) + \sqrt{\frac{1}{L}}\frac{cosh(\pi(2n-1)s/2L)}{sinh(\pi(2n-1)/4)} \; n\ even \end{cases} \tag{21}$$

$$\Lambda_n \approx \frac{3k_BT\lambda\pi^4(2n-1)^4}{32L^4} \tag{22}$$

$$\sum_{n\neq m} \frac{H_{nm}^2\Lambda_m}{H_{nn}\Lambda_n - H_{mm}\Lambda_m} \approx \frac{12\sqrt{6}}{n\sqrt{\pi}\left(3+A-12log(\frac{L}{an\pi})\right)} \tag{23}$$

$$\sum_{n\neq m} \psi_m(L/2)\frac{H_{nm}\Lambda_m}{H_{nn}\Lambda_n - H_{mm}\Lambda_m} \approx \frac{2}{\sqrt{L}}\left(\frac{1}{3+A-12log\left(\frac{L}{an\pi}\right)}\right) \tag{24}$$

where constant $A = 18\gamma - 2(6\pi)^{1/2} + 6E_1(3/2) + 6log(6) \approx 13.06$

Hinczweski *et al.* approximations, described above, take in full account of the hydrodynamic interactions of the polymers and are successfully validated on experiments performed on various lengths of dsDNA [13–15]. We further implemented the theoretical framework developed by Hinczweski *et al.* for finding the $\lambda$ in the different conformations of EEA1. We used $a$ and $L$ as the fixed parameters obtained from the partially solved crystal structure of EEA1 ($PDB : 1JOC$) ($a = 1nm$) and electron microscopy rotary shadowing data from Murray *et al.* ($L = 220nm$) [16]. By fitting Eq. 6 to the earlier obtained $\langle r^2(\tau)\rangle$ through FCS measurements, we obtained the persistence length ($\lambda$) for EEA1 in various conformational states.

##### *4. Bootstrapping*

The fraction of EEA1 molecules either in collapsed or extended state varies progressively over time. Fig. 2.a describes the average local exponent $\alpha$ for the mixed population of EEA1 conformations. To obtain the distributions of the population in collapsed and extended state bootstrapping was performed on the dcFCCS data. Bootstrapping is performed on the set of 180 curves. Randomly, 20 of these curves are selected and their mean at each lag-time ($\tau$) calculated to obtain a single curve. The mean dcFCCS curve is then used to calculate $MSD(\tau)$ using Eq. 1. The process is repeated to obtain $10^5$ dcFCCS mean curves and resulting $MSD(\tau)$. The data is then further analysed to obtain the minima of the local exponent $\alpha_{min}$ and the persistence length $\lambda$ by applying approximations derived by Hinczweski *et al.* [13].

To observe the recovery from flexible to extended state, we first computed the local exponent from the $MSD(\tau)$ over time using a moving window of 15 minutes (Fig. 2a). We performed bootstrapping to obtain distributions of $\alpha_{min}$ (sample size, S= $10^5$). $\alpha_{min}$ is obtained by a custom written computer code that finds the minimum of the local exponent ($\alpha(\tau)$) and then takes the average of previous and next 3 points for the lag-times. Fig. 2b shows the $\alpha_{min}$ mean ($\bar{x} = \frac{1}{S}\sum_{i=1}^{S} x_i$) with open circles and standard error of the mean ($SDE_\alpha$) with error bars calculated as follows:

$$SDE_\alpha = \begin{cases} \sqrt{\frac{1}{S_h \times 180}\sum_{i=1}^{S_h-1}(x_i-\bar{x})^2}, \forall x_i > \bar{x} \\ \\ \sqrt{\frac{1}{S_l \times 180}\sum_{i=1}^{S_l-1}(x_i-\bar{x})^2}, \forall x_i \leq \bar{x} \end{cases} \tag{25}$$

where $S_h$ are the number of elements $\forall x_i > \bar{x}$, $S_l$ are the number of elements $\forall x_i \leq \bar{x}$ and

$S_h + S_l = S$. For EEA1 $\alpha_{min} = 0.72$, indicating that the protein is in an extended state. Upon addition of Rab5(GTP) $\alpha_{min} = 0.66$, and then recovers overtime to $\alpha_{min} = 0.75$. Similarly, in the second round of Rab5(GTP) addition, the $\alpha_{min}$ reduces to 0.67, and over time recovers back to 3/4 scaling local exponent (solid blue line). This confirms that EEA1 population can perform multiple cycles of collapse and extension.

Later, the bootstrapping strategy is used to calculate the distribution of persistence length of EEA1 by applying Hinczweski's approximations. We obtained the persistence length ($\lambda$) change over the two cycles of EEA1 collapse and re-extension (Fig. 2c). Next, we compare the distributions of the $\lambda$. The open circle represent the peak value of the distribution and the error bar represents standard error of the peak calculated by Eq.26 as follows:

$$SDE_\lambda = \begin{cases} \sqrt{\frac{1}{S_h \times 180} \sum_{i=1}^{S_h - 1} (x_i - x_p)^2}, \forall x_i > x_p \\ \\ \sqrt{\frac{1}{S_l \times 180} \sum_{i=1}^{S_l - 1} (x_i - x_p)^2}, \forall x_i \leq x_p \end{cases} \tag{26}$$

where $x_p$ is peak value of distribution, $S_h$ is the number of elements $\forall\ x_i > x_p$ and $S_l$ is the number of elements $\forall\ x_i \leq x_p$. For EEA1 a single peak was observed at $\approx 170nm$. Immediately after each addition of Rab5(GTP) we observed single peaks at $\approx 67nm$ and $\approx 69nm$ for cycle1 and cycle2 respectively. However, over time there is an onset of a second peak revealing the existence of a mixed population of collapsed and extended EEA1 molecules (see Extended Data Fig. 6). Hence, we take the value of the second peak to show the population recovery and calculate $SDE_\lambda$ from the second peak. The second peak appears at $\approx 190nm$ for both cycle1 and cycle2, which confirms that the population recovers to the original extended state. In the same distribution, a peak at $\approx 70nm$ persistent across all time-points, suggests that exists mixed populations for a long time.

##### 5. *Preference of FCS over dcFCCS*

To access the faster time-scales we make use of dcFCCS as it avoids the overlap of time-scales coming from fluctuations of the polymer end and the photophysical events such as triplet state and detector noise. Extended Data Fig. 4 shows the comparison between single color FCS and dcFCCS, the detector noise is marked with an ellipse and the triplet state with a rectangular region for FCS (green and red channels). In contrast, dcFCCS doesn't show detector noise nor triplet states, which allows access to the faster time-scales of the polymer's internal dynamics.

##### F. Experiments to probe EEA1 flexibility in different states

Experiments were carried out in a 384-well glass bottom plate with $175 \pm 15\mu$m glass thickness. The plate was sealed with aluminium foil to prevent drying. All measurements were performed in the same setup, where the confocal beam, pinhole alignment, laser power and the collar adjustments were kept constant across all experiments.

The experiment was performed as follows: $50\mu$l of EEA1 ($100nM$) were added to a well and the confocal beams were focused $20\mu$m above the glass surface to perform dcFCCS. 240 correlation curves of 30 seconds each were recorded for 2 hours. We observed that EEA1 sticks to the glass surface. Thus, the first 30min recordings for EEA1 alone were not analysed. They were used to stabilize the system by allowing EEA1 to stick to the surface of the well and bleach the area used for measurements. Further, we added two rounds of $2\mu M$ Rab5(GTP) to the solution and dcFCCS was evaluated for 2.5 hours after each addition. We evaluated the EEA1 dynamics over long time to determine if EEA1 can recover from its collapsed state (reversibility) and also, if the resulting collapse-extension cycle could be repeated (recyclability) (See Fig. 2).

As a negative control we evaluated the effect of having EEA1 mixed with inactive Rab5(GDP). $50\mu l$ of EEA1 ($100nM$) were added to another well and again performed dcFCCS for next 2 hours (240 curves). In the next step, we added $2\mu M$ Rab5(GDP) to the same well and perfomed dcFCCS measurements for 1.5 hours. As shown in Extended Data Fig. 5, there is no reduction in the $\alpha_{min}$ for EEA1 in presence of Rab5(GDP) in comparison to EEA1 alone. Rather, we see a slight increase in the $\alpha_{min}$, which we suspect is because of non-specific interactions between Rab5(GDP) and EEA1.

### III. POLYMER MODEL FOR THE EEA1-RAB5 TWO COMPONENT MOLECULAR MOTOR

#### A. Free energy, elastic energy and conformational entropy in the Blundell-Terentjew model

We use a coarse-grained model to capture the essential features of the EEA1-Rab5 system. Here we model EEA1 as a semiflexible polymer with a contour length, $L$, and an effective persistence length, $\lambda_h$. The interaction with active Rab5 at the amino-terminal tip of EEA1 triggers a global stiffness transition and results in a functional EEA1:Rab5 complex with a much reduced effective persistence length $\lambda_l$, where $\lambda_l < \lambda_h \sim L$. The system works under isothermal conditions.

To a first approximation, we assume that the contour length of EEA1, $L$, does not change significantly between the two states, which are EEA1 unbound and EEA1 bound to Rab5. Previously, a contour length of $L = (222 \pm 26)\,\mathrm{nm}$ was reported for EEA1 alone, and $L = (195 \pm 26)\,\mathrm{nm}$ for the condition of EEA1 plus Rab5(GTP) [16]. For the idealized model presented in this study, we chose the rounded average value between these two contour length measurements, i.e. $L = 210\,\mathrm{nm}$ as our fixed contour length for all calculations. Exploring the role of a contour length change of up to 10 % did not lead to qualitatively different model behaviour, indicating that the dominant effect can be explained by the much larger change of the effective persistence length.

The free energy in the Blundell-Terentjew model of an inextensible semiflexible polymer is given by [17]:

$$A(x, \kappa) = \frac{\kappa \pi^2}{2L}\left(1 - x^2\right) + \frac{2\left(k_{\mathrm{B}}T\right)^2 L}{\pi \kappa \left(1 - x^2\right)} \tag{27}$$

with contour length $L$, relative extension $x = r/L$, and bending modulus $\kappa = \lambda \cdot k_{\mathrm{B}}T$, which relates thermal energy to the persistence length $\lambda$.

If we substitute the expression for $\kappa$ and separate the thermal energy, we get:

$$a(x, \lambda) = A/k_{\mathrm{B}}T = \frac{\lambda \pi^2}{2L}\left(1 - x^2\right) + T \times \frac{2L}{\pi\, T\, \lambda\, (1 - x^2)} \tag{28}$$

Following Blundell & Terentjew, we identify the first term with the internal elastic bending energy

$$u(x,\lambda) = U/k_{\mathrm{B}}T = \frac{\lambda\pi^2}{2L}\left(1-x^2\right), \tag{29}$$

and the second term with the conformational entropy

$$s(x,\lambda) = S/k_{\mathrm{B}}T = -\frac{2L}{\pi\, T\, \lambda\, (1-x^2)}. \tag{30}$$

Using these two term we obtain the familiar relation:

$$a = u - T \times s \tag{31}$$

In this model, the equilibrium extension at zero force is given by $r_0(\lambda) = L\sqrt{1 - \frac{2L}{\pi^{3/2}\lambda}}$ and characterises the minimum of the corresponding free energy. The critical persistence length is given by $\lambda_c = 2\pi^{-3/2}L$ and indicates the value below which the root in the expression for the equilibrium extension has a complex solution and the equilibrium extension is set to zero, *i.e.* $r_0(\lambda < \lambda_c) = 0$. Using these general expressions, we define the *capture distance* as the equilibrium extension of the unbound EEA1 with the higher persistence length $\lambda_{\mathrm{h}}$ as $r_{\mathrm{capture}} = r_0(\lambda_{\mathrm{h}})$. Likewise, we define the *release distance* as the equilibrium extension of the EEA1:Rab5 complex with the lower persistence length $\lambda_{\mathrm{l}}$, as $r_{\mathrm{release}} = r_0(\lambda_{\mathrm{l}})$. In all these discussions we assume that the contour length, $L$, stays constant. Interestingly, at mechanical equilibrium one obtains the values $u(r_0) = \sqrt{\pi}$ and $s(r_0) = -\sqrt{\pi}/T$ for the bending energy and conformational entropy, leading to equilibrium free energy value $a(r_0) = 2\sqrt{\pi}$.

Furthermore, the force can be obtained from the free energy as $F(r,\lambda) = -\partial_r A(r,\lambda)$:

$$F(r,\lambda) = \frac{4\, r\, k_{\mathrm{B}}T}{\pi\lambda\, L\, (1-r^2/L^2)^2} - \frac{\pi^2 r\, \lambda\, k_{\mathrm{B}}T}{L^3} \tag{32}$$

The mechanical work performed during the entropic collapse or the recovery step is obtained by integrating the force between the capture and release extensions, which we defined above.

$$W(r,\lambda) = \int_{r_{\mathrm{capture}}}^{r_{\mathrm{release}}} \mathrm{d}r F(r,\lambda) \tag{33}$$

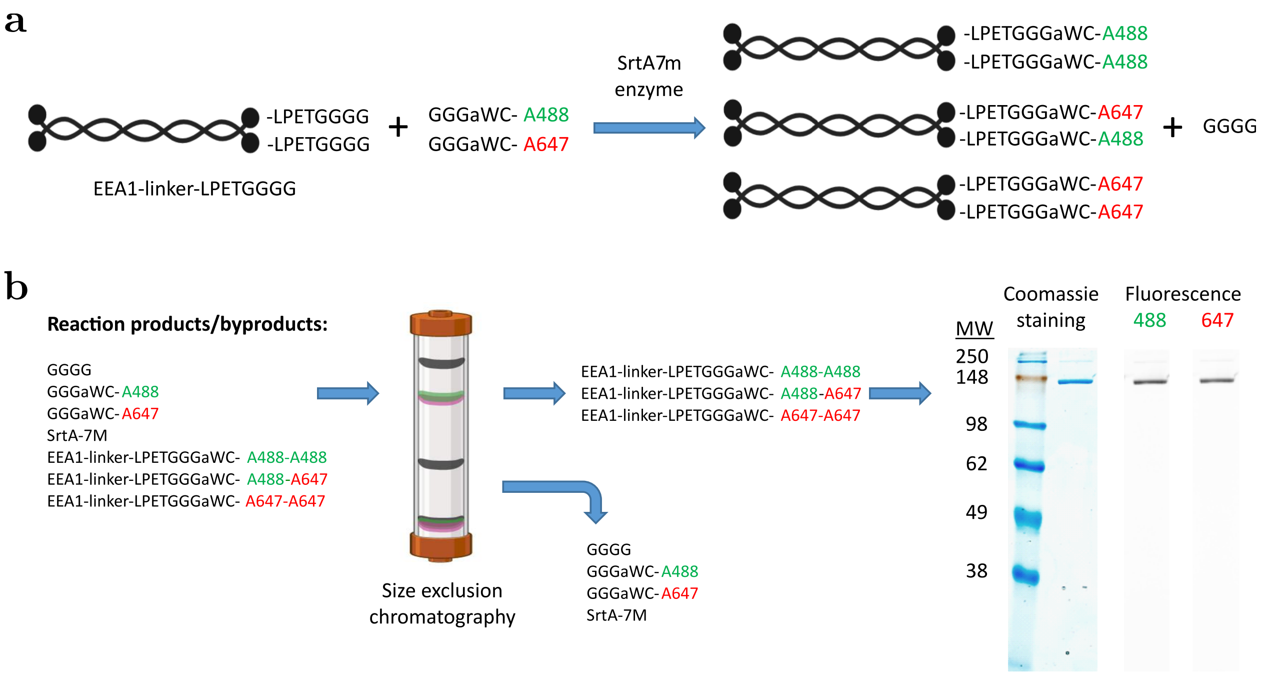


**Extended Data Fig.** 1. **EEA1 dual colour labelling and purification (a)** EEA1 C-terminal labelling with Alexa488 and Alexa647 using SrtA7m enzyme. Labelling to each EEA1 monomer is independent, resulting in EEA1 with two Alexa488, two Alexa647 and the desired configuration of one fluorophore of each type. **(b)** Purification of dual-labelled EEA1 by size exclusion chromatography to remove "GGGG" reaction by-product, excess dye, and the enzyme. Purified protein was assessed by SDS-PAGE followed by laser scanning (488 and 647nm) and coomassie staining. A band is observed in all cases indicating successful labelling.

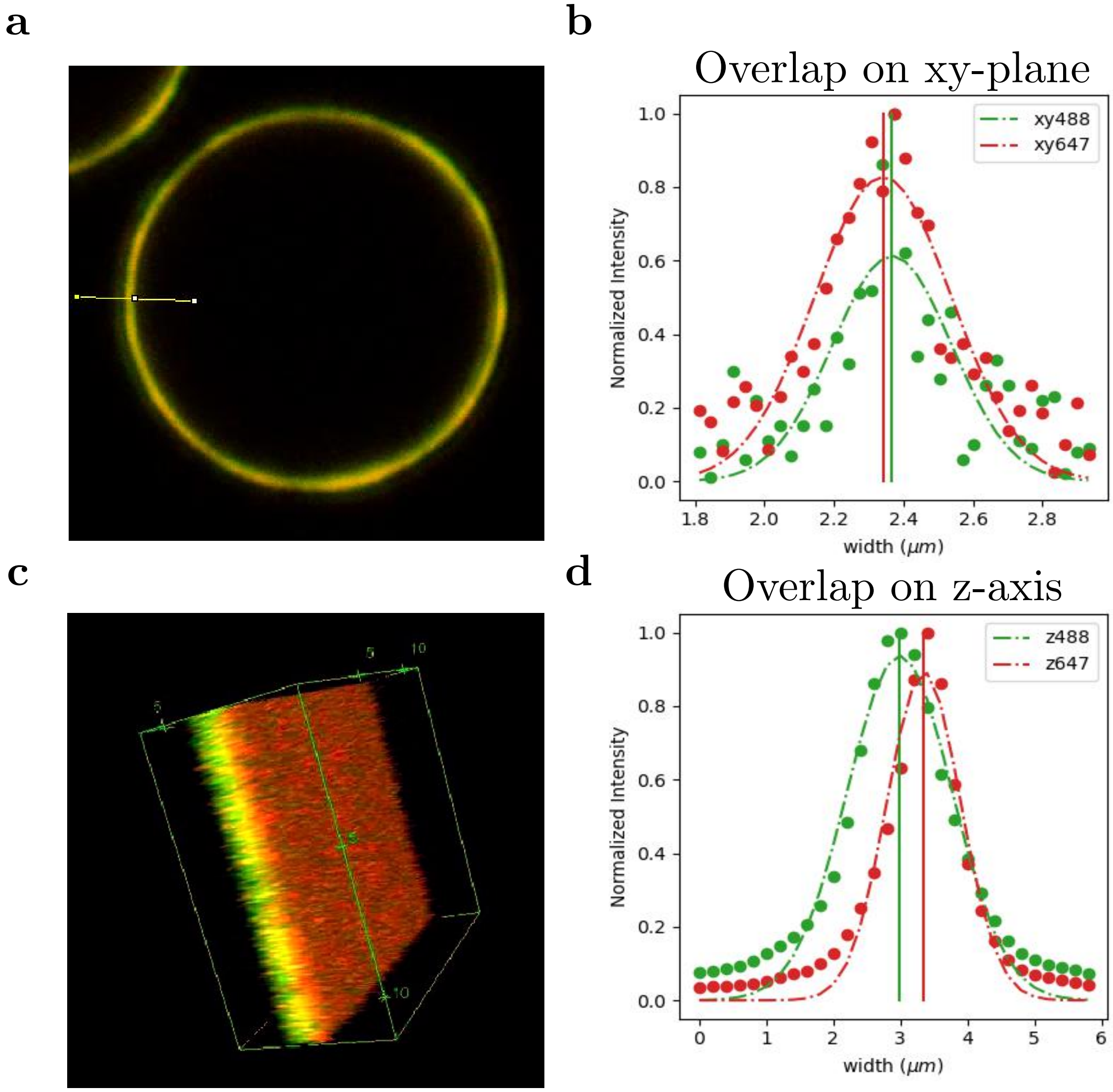


**Extended Data Fig.** 2. **Confocal volume overlap in lateral** $(X, Y)$ **and axial** $(Z)$ **directions** Images show the merge of DIO(green) and DID(red) channels. **(a)** Confocal image of a $10\mu m$ MCB at the equatorial plane, which is used to calculate the overlap between channels. Yellow horizontal line (X-axis) indicates the region used to evaluate the intensity profile of each channel. Normalized intensities are plotted in **(b)**. Obtained a peak difference of $23nm$ and 84% area overlap. **(c)** 3D reconstruction from a confocal Z-stack of a SLB. Used to calculate the overlap between channels in the Z-axis. Normalized mean intensities of each plane are plotted in **(d)**. Obtained a peak difference of $367nm$ and 65% area overlap.

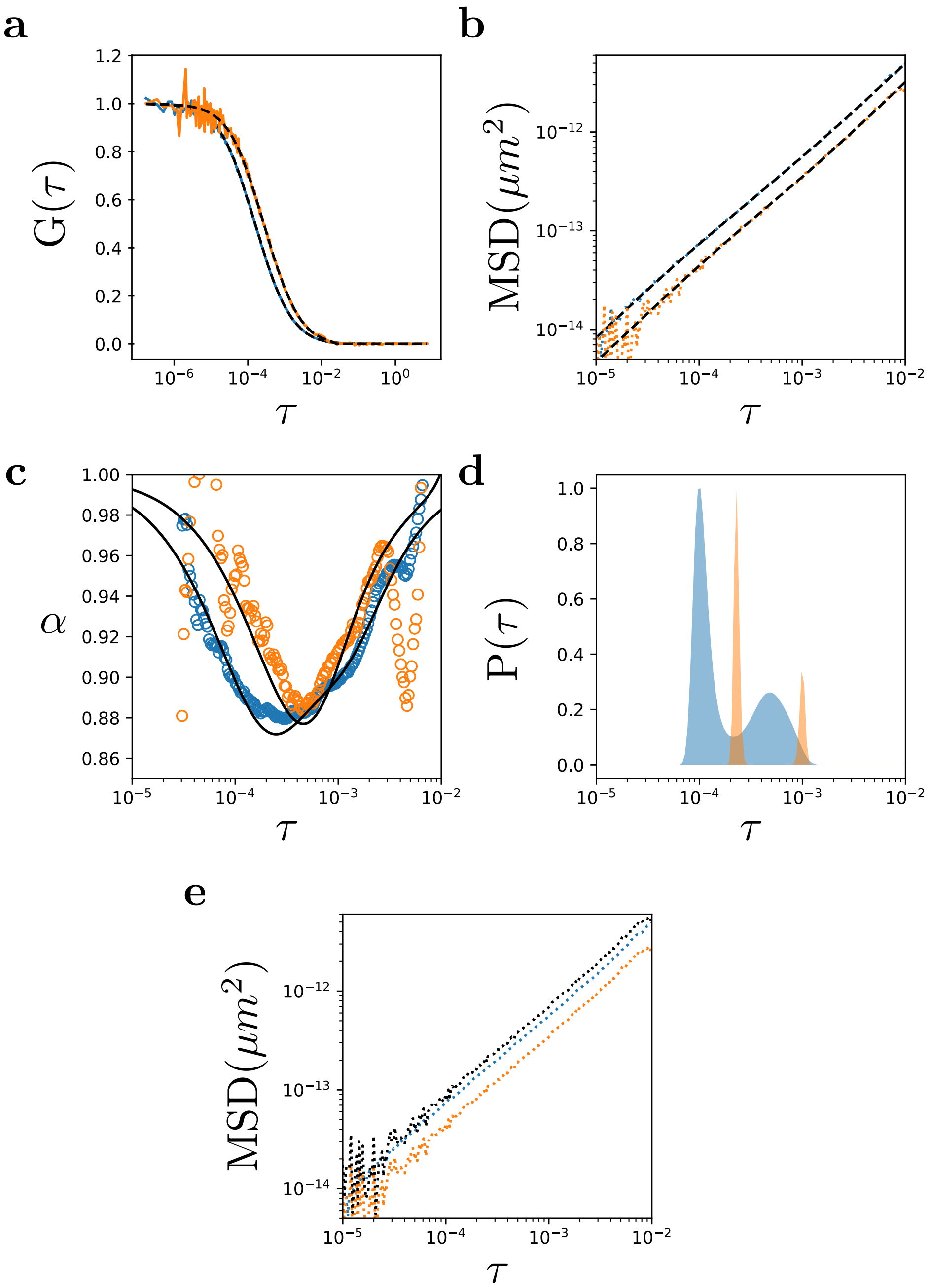


**Extended Data Fig.** 3. **Calibration of the overlapping confocal volumes.** **(a)** Shows the auto-correlation(blue) and cross-correlation(orange) for FCS and dcFCCS. **(b)** Shows the mean-square displacement calculated by using the confocal width of FCS for both correlation curves. **(c)** Plots the local slopes $\alpha(\tau)$ of the two $MSD(\tau)$ curves. **(d)** Probabilities of time-scales present in each correlation curve are plotted. **(e)** With the corrected confocal volume of the dcFCCS the adjusted $MSD(\tau)$ is plotted with black dotted curve, which sits next to FCS(blue)

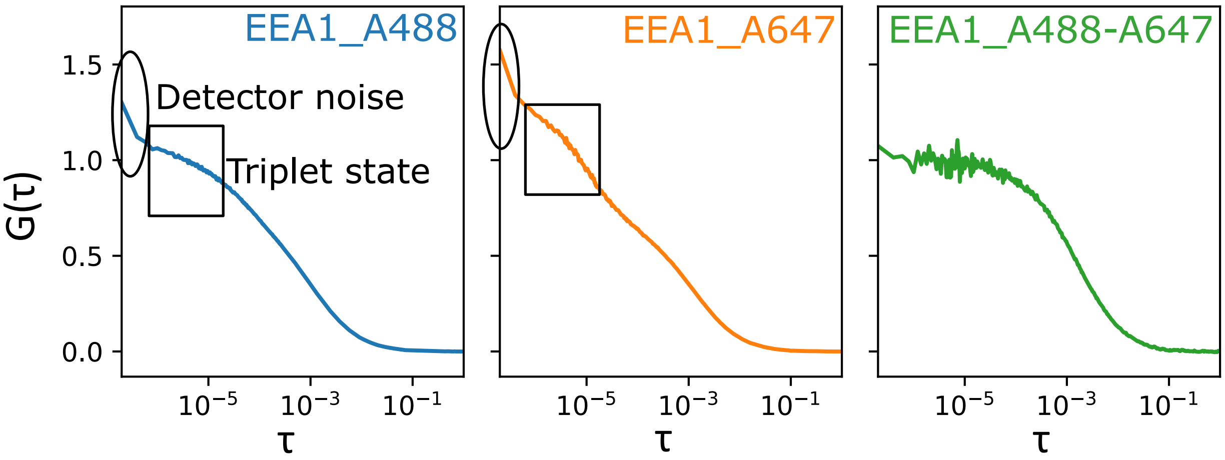


**Extended Data Fig.** 4. **Highlighting the difference between FCS and dcFCCS** Shows the auto-correlation (FCS) for EEA1_A488(blue) and EEA1_A647(orange); and cross-correlation (dcFCCS) for EEA1_A488_A647(green). The artifacts caused by detector noise and triplet state of the fluorophores are highlighted with an ellipse and a rectangular region, respectively.

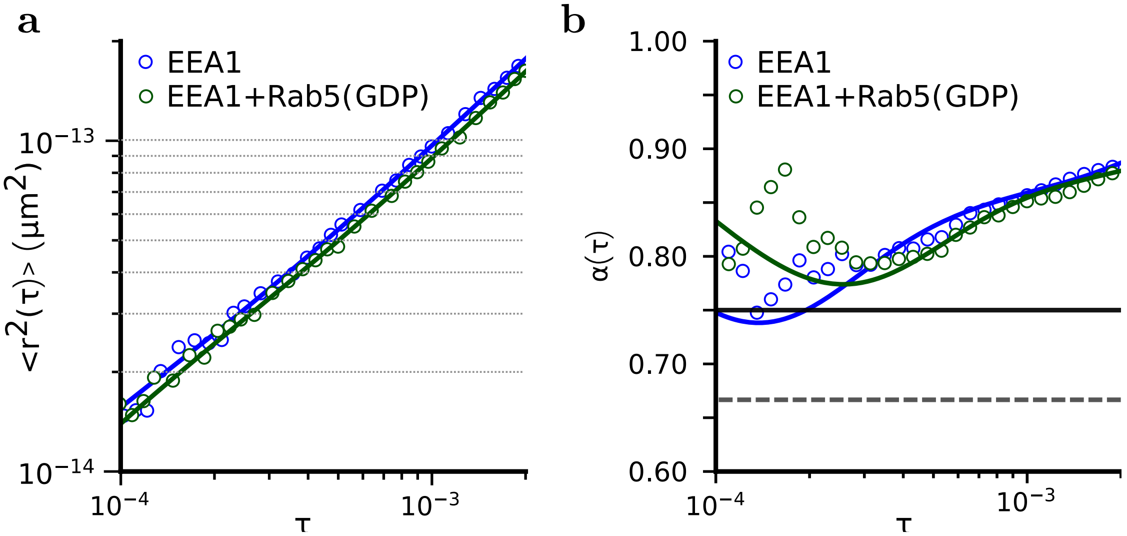


**Extended Data Fig.** 5. **Effects of non-specific interactions of EEA1 with Rab5(GDP)** The dynamics in solution for EEA1 alone (blue) and when mixed with Rab5(GDP) (green) are quantified by the **(a)** mean square displacement (MSD) plotted over a lag time $\tau$ and **(b)** the local scaling exponent $\alpha$ of the MSD. **(a)** Shows the dynamics of EEA1 being affected by the non-specific interaction with Rab5(GDP). **(b)** Shows an slight increase in the local scaling exponent $\alpha$ upon addition of Rab5(GDP), reflecting a change in EEA1 internal dynamics.

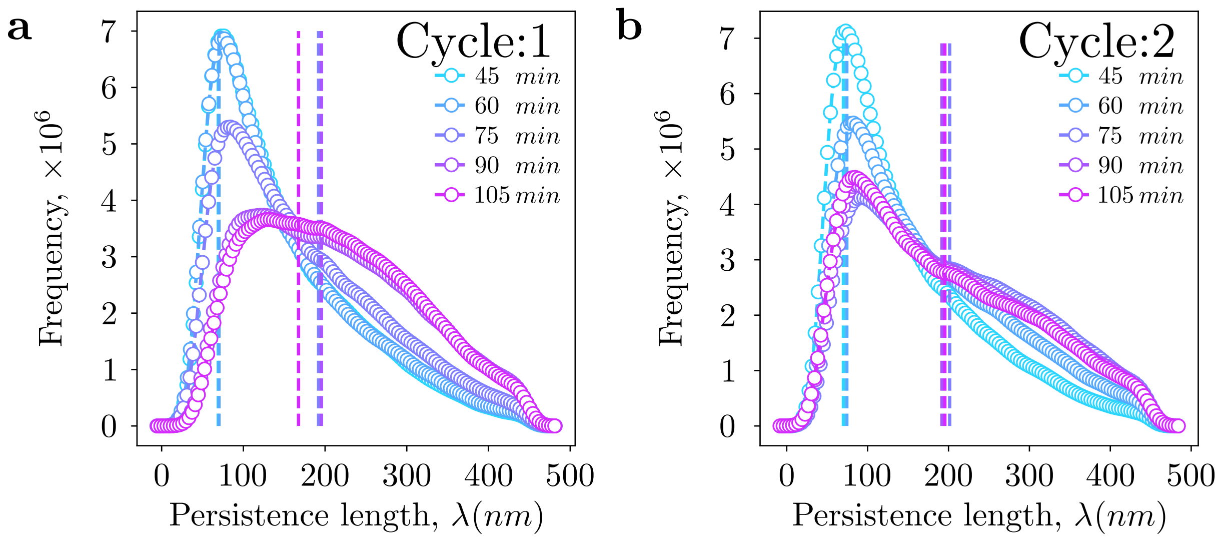


**Extended Data Fig.** 6. **Persistence length distribution at different time-points in the EEA1 extension-collapse cycles.** **(a)** and **(b)** show change in the distribution of persistence length during the first and second cycle respectively. The change in distribution over time is represented with an spectra from cyan to magenta, *i.e.* from 45 $min$ to 105 $min$ for both the cycles. The identified peaks of the distributions are represented with dashed lines.

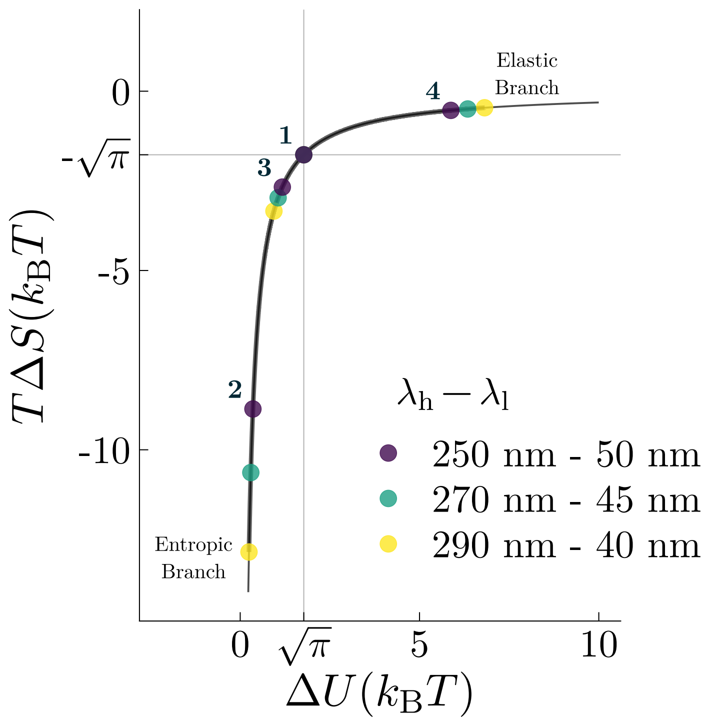


**Extended Data Fig.** 7. **Disturbing the balance between entropic and enthalpic contributions over the cycle.** During a full work cycle of a semiflexible polymer motor, the balance between conformational entropy and bending energy traverses along a universal curve. First, the equilibrium of the free polymer is characterised by the exact balance between enthalpic and entropic contributions (state 1). Second, the softening at the constant capture extension drives the two-component system deep into the entropic branch (state 2). From here, the complex climbs back through entropic collapse to state 3. During the second half of the cycle, the release of the GTPase causes a sudden increase in the rigidity at the collapsed extension. This leads to a movement into the elastic branch (state 4). Finally, the free polymer recovers its balance and equilibrium at state 1. In the asymptotic limits of zero temperature or infinitely stiff rods, the entropic contribution is zero. Similarly, for perfectly flexible polymers the contribution from the bending energy also approaches zero. State colors indicate different pairings of the two flexibility values. The larger the difference between the high and low effective persistence length, the deeper the system moves into the branches. Cycle states are only indicated for $\lambda_h = 250\,\text{nm}$ and $\lambda_l = 50\,\text{nm}$.

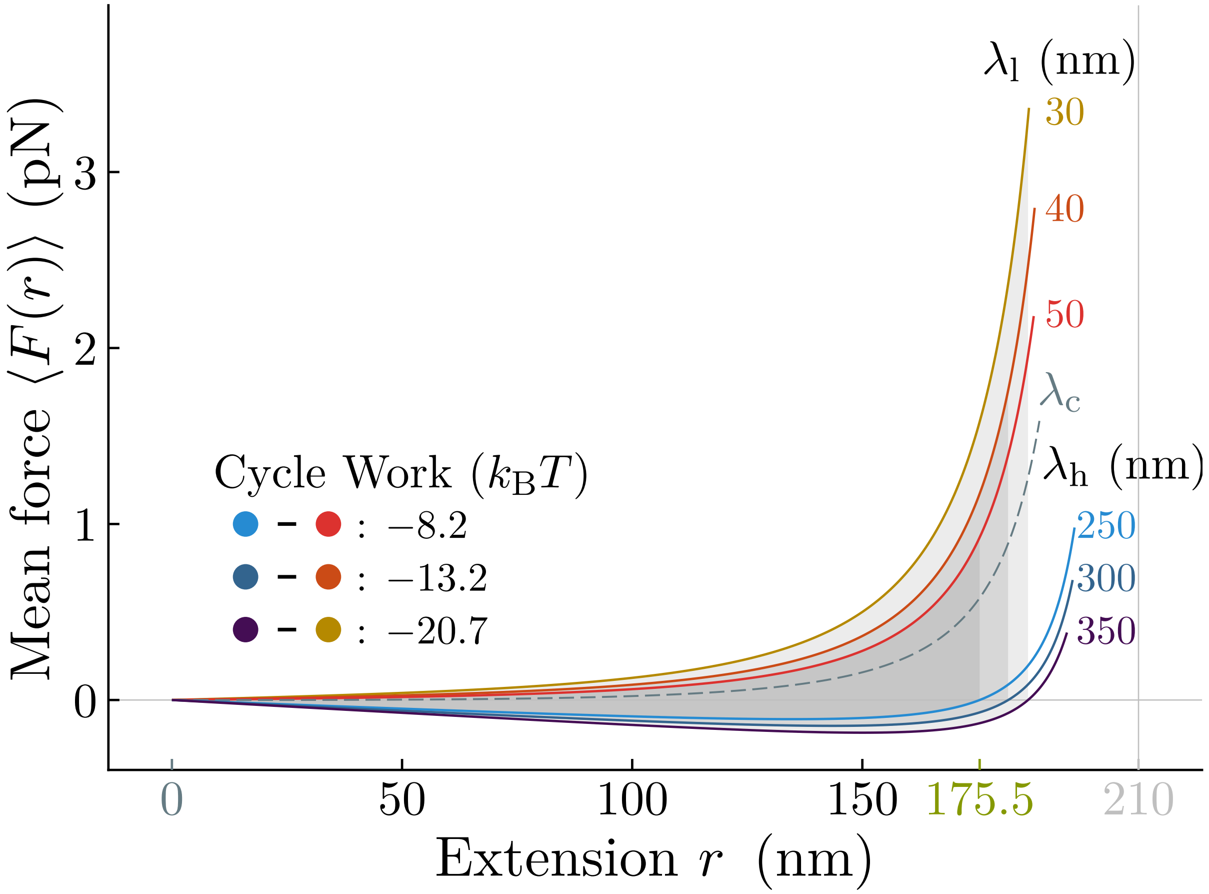


**Extended Data Fig.** 8. **The maximum extractable work in a Sterling-like polymer engine depends on the pairing of stiffnesses.** Different effective persistence lengths result in different magnitudes of the work performed within a cycle. The two force-extension curves enclose successively larger areas with a larger separation between the two states. To illustrate the effect, three pairs with successively larger differences between their higher and lower persistence lengths are illustrated: $\lambda_h = 250\,\mathrm{nm} - \lambda_l = 50\,\mathrm{nm}$ (blue-red), $\lambda_h = 300\,\mathrm{nm} - \lambda_l = 40\,\mathrm{nm}$ (petrol-orange), $\lambda_h = 350\,\mathrm{nm} - \lambda_l = 30\,\mathrm{nm}$ (dark purple-yellow).

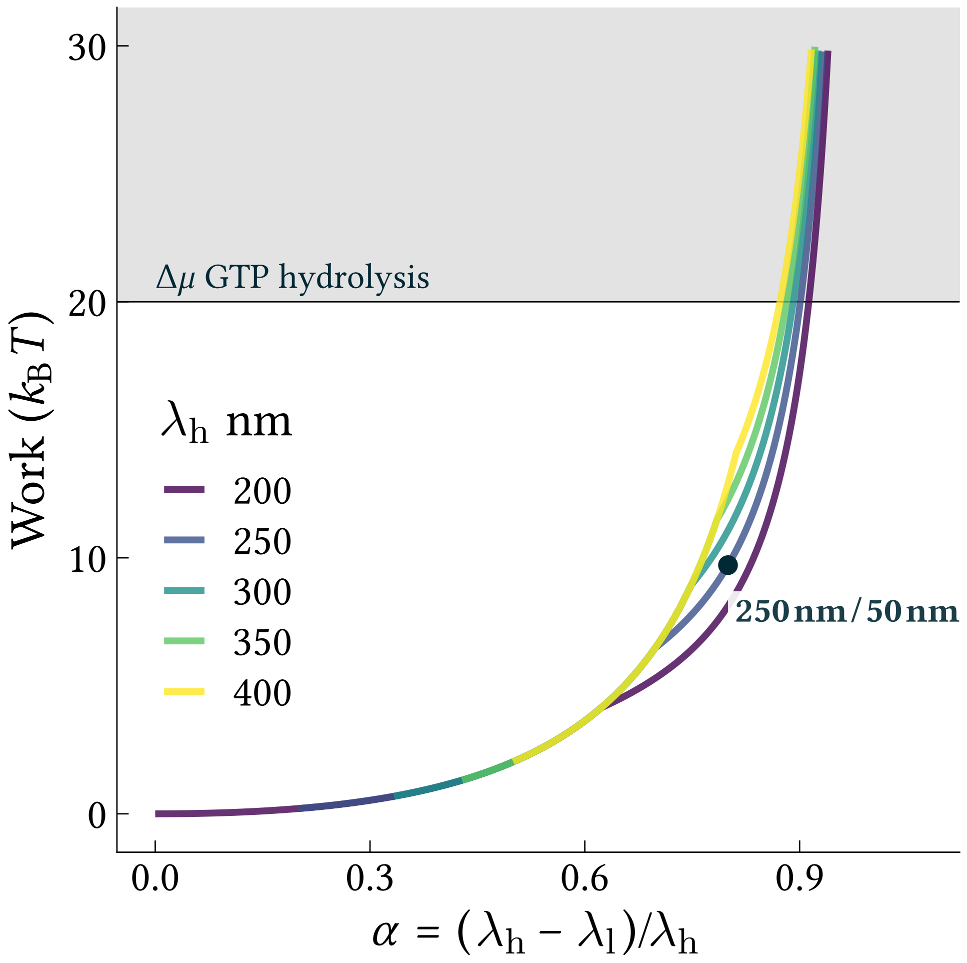


**Extended Data Fig.** 9. **Efficiency of an idealized WLC polymer engine.** The amount of mechanical work that can be extracted during a full mechano-chemical cycle depends on the difference between the higher and the lower effective persistence lengths of the semiflexible tether. Here, the two quantities are combined into a dimensionless quantity, $\alpha = (\lambda_\mathrm{h} - \lambda_\mathrm{l})/\lambda_\mathrm{h}$, ranging from 0, if $\lambda_\mathrm{l} = \lambda_\mathrm{h}$, to 1, if $\lambda_\mathrm{l} = 0\,\mathrm{nm}$. Different colours indicate different values of the unbound value, $\lambda_\mathrm{h}$. An appreciable amount of mechanical work ($> 5\,k_\mathrm{B}T$) requires at least a value of $\alpha > 0.5$. Indicated is also a comparison to the available energy budget from GTP hydrolysis, assumed here for illustrative purposes to be $20\,k_\mathrm{B}T$. Note that in a similar plot for an ideal gas the canonical Carnot efficiency would be a strait line, which highlights the non-linear nature of polymer engines.
